# Disentangling microbial networks across pelagic zones in the global ocean

**DOI:** 10.1101/2021.07.12.451729

**Authors:** Ina M. Deutschmann, Erwan Delage, Caterina R. Giner, Marta Sebastián, Julie Poulain, Javier Arístegui, Carlos M. Duarte, Silvia G. Acinas, Ramon Massana, Josep M. Gasol, Damien Eveillard, Samuel Chaffron, Ramiro Logares

## Abstract

Microbial interactions underpin ocean ecosystem function, but they remain barely known. Multiple studies have analyzed microbial interactions using static association networks based on omics data, yet microbial interactions are dynamic and can change across spatiotemporal scales. Understanding the dynamics of microbial interactions is needed for a better comprehension of ocean ecosystems. Here, we explored associations between archaea, bacteria, and picoeukaryotes along the water column, from the surface to the deep ocean, across the northern subtropical to the southern temperate ocean and the Mediterranean Sea by defining sample-specific subnetworks, which allowed us to examine changes in microbial associations across space. We found that associations tend to change with depth as well as with geographical scale, with a few associations being global (i.e., present across regions within the same depth layer) and 11-36% being regional within specific water layers. The lowest fraction of global associations was found in the bathypelagic zone, while associations restricted to certain regions increased with depth. The majority of associations observed in surface waters disappeared with depth, suggesting that surface ocean associations are not transferred to the deep sea, despite microbial sinking. Altogether, our results suggest that microbial associations have highly heterogeneous distributions in the horizontal and vertical dimensions of the ocean and that such distributions do not mirror taxonomic distributions. Our work contributes to better understand the dynamics of microbial interactions in the global ocean, which is urgently needed in a context of global change.

## INTRODUCTION

Microorganisms play fundamental roles in ocean ecosystem functioning and global biogeochemical cycles (1–3). The main processes shaping microbial community composition are selection, dispersal, and drift (4). Selection exerted via environmental heterogeneity and biotic interactions is essential in structuring the ocean microbiome (5), leading to heterogeneities in community composition that can reflect those found in the ocean, normally related with temperature, light, pressure, nutrients, and salinity. Global-scale studies of the surface ocean reported strong associations between microbial community composition and temperature (5–8). Marked changes in microbial communities with depth have also been reported (9–14), reflecting the steep vertical gradients in light, temperature, nutrients and pressure.

Prokaryotes (bacteria and archaea) and unicellular eukaryotes are fundamentally different in terms of ecological roles, functional versatility, and evolutionary history (15) and are connected through biogeochemical and food web interaction networks (16,17). Still, our knowledge about their ecological interactions remains limited, even though these interactions sustain marine food webs and contribute to nutrient recycling in the oceans (3,18). Microbial interactions are very difficult to resolve experimentally, mainly because most microorganisms are hard to cultivate (19,20) and synthetic laboratory communities are unlikely to mirror the complexity of wild communities. However, association networks inferred from omics data have the potential to unravel microbial interactions.

Microbial association networks are normally based on abundance data, representing putative ecological interactions that need to be confirmed via laboratory experiments. Yet, association networks are one of the best available tools to start addressing the huge complexity of microbial interactions. Association networks can provide a general overview of the potential microbial interactions in the ocean aggregated over a given period of time (9,10,21–25) or through space (26–28). Previous work investigated marine microbial associations within and across depths. For example, prokaryotic associations were investigated in the San Pedro Channel, off the coast of Los Angeles, California, covering the water column from the surface (5 m) to the seafloor (890 m) (9,10). Furthermore, a global survey from the TARA Oceans expedition investigated planktonic associations between a range of organismal size fractions in the epipelagic zone (26), from pole to pole (28). However, these studies did not include the bathypelagic realm below 1000 m depth, which represents the largest microbial habitat in the biosphere (29).

Most studies so far have investigated microbial associations in the ocean using static networks determined from spatially distributed samples, which capture global, regional and local associations in a single network. Furthermore, given that global-ocean expeditions collect samples over several months, networks must include some temporal associations, yet disentangling them from spatial associations is challenging. Spatially widespread or global associations may be part of the core microbiome defined as the set of interacting microbes essential for the functioning of the ocean ecosystem (30). Core associations may be detected by constructing a single network from numerous locations and identifying the most significant and strongest associations (31). In turn, regional or local associations may reflect interactions occurring in specific locations due to taxa distributions resulting from abiotic or biotic environmental selection, or dispersal limitation. Regional networks could also contribute to determine associations that are stable (i.e., two partners always together) or variable (one partner able to interact with multiple partners across locations). The fraction of regional associations may be determined by excluding all samples belonging to one region, recomputing network inference with the reduced dataset, and examining which associations are missing (26). Alternatively, regional networks are computed considering samples belonging to the regions, allowing to determine both global and regional associations (32) by investigating which edges are common and which are unique.

Regional networks, however, require a high number of samples per delineated zone and these may not be available due to logistic or budgetary limitations. Recent approaches circumvent this limitation by deriving sample-specific subnetworks from a single static, i.e., all-sample network, which allows quantifying association recurrence over spatiotemporal scales (28,33). Here, we adjusted the approach to determine global and regional associations along vertical and horizontal pelagic ocean scales, which allowed us determining a biogeography of marine microbial associations. We analyzed associations between archaea, bacteria, and picoeukaryotes using a unique dataset including 397 samples covering the water column, from surface to deep waters, in the Mediterranean Sea (hereafter MS) and five ocean basins: North and South Atlantic Ocean, North and South Pacific Ocean, and Indian Ocean (hereafter NAO, SAO, NPO, SPO, and IO) (Figure 1). Our exploration of the variation of subnetworks across regions and depths allowed us to determine widespread associations as well as local associations that seem to be only present in specific locations or depths.

**Figure 1:**
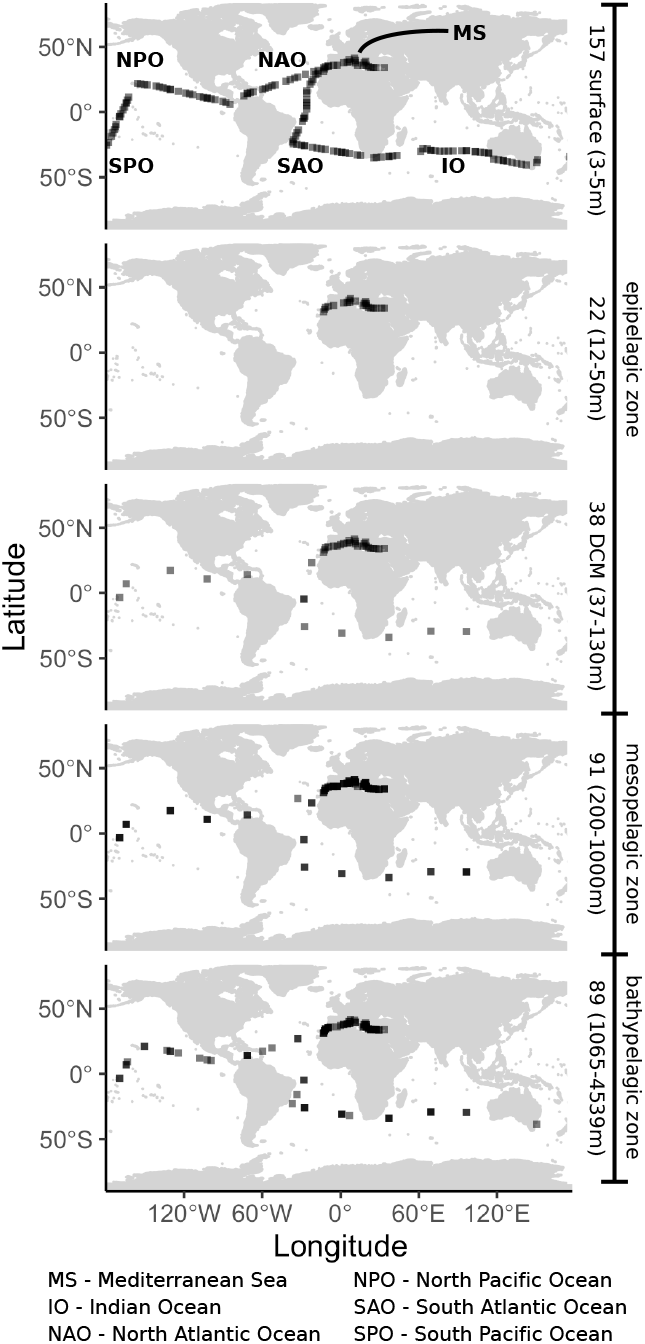
Sampling scheme. Location, number, and depth range of samples from the epipelagic zone including surface and DCM layers, the mesopelagic zone, and the bathypelagic zone from the global tropical and subtropical ocean, and the Mediterranean Sea.

## RESULTS

### Network architecture changed along the water column

Microbial dispersal as well as vertical and horizontal environmental heterogeneity are expected to affect network topologies. Yet, we have a limited understanding on how much marine microbial networks change due to these processes, and analyzing the topology of subnetworks from specific ocean regions and depths is a first step to address this issue. We generated 397 sample-specific subnetworks and compared them across the regions and depth layers using eight network metrics (see Methods). We found that network metrics change along the water column (Supplementary Figure 1). As a general trend, subnetworks from deeper zones were more clustered (transitivity), had higher average path length, featured stronger associations (average positive association scores), and lower assortativity (based on degree) compared to those in surface waters. Most subnetworks from the Deep Chlorophyll Maximum (DCM) and bathypelagic zones had the highest edge density, i.e., highest node connectivity. In contrast, in the MS, the surface subnetworks had the highest node connectivity (Supplementary Figure 1).

### Only a few global associations

We computed the spatial recurrence, i.e., prevalence, of each association as the fraction of subnetworks in which a given association was present across all 397 subnetworks (Figure 2A) and within each region-depth-layer combination (Figure 2B). The global ocean surface layer (contributing 40% of the samples) had more associations compared to the other depths (Figure 2B). Remarkably, 14,971 out of 18,234 (82.1%) surface ocean associations detected in the basins were absent in the MS. In turn, the number of surface associations was similar across the five ocean basins (Figure 2B).

**Figure 2:**
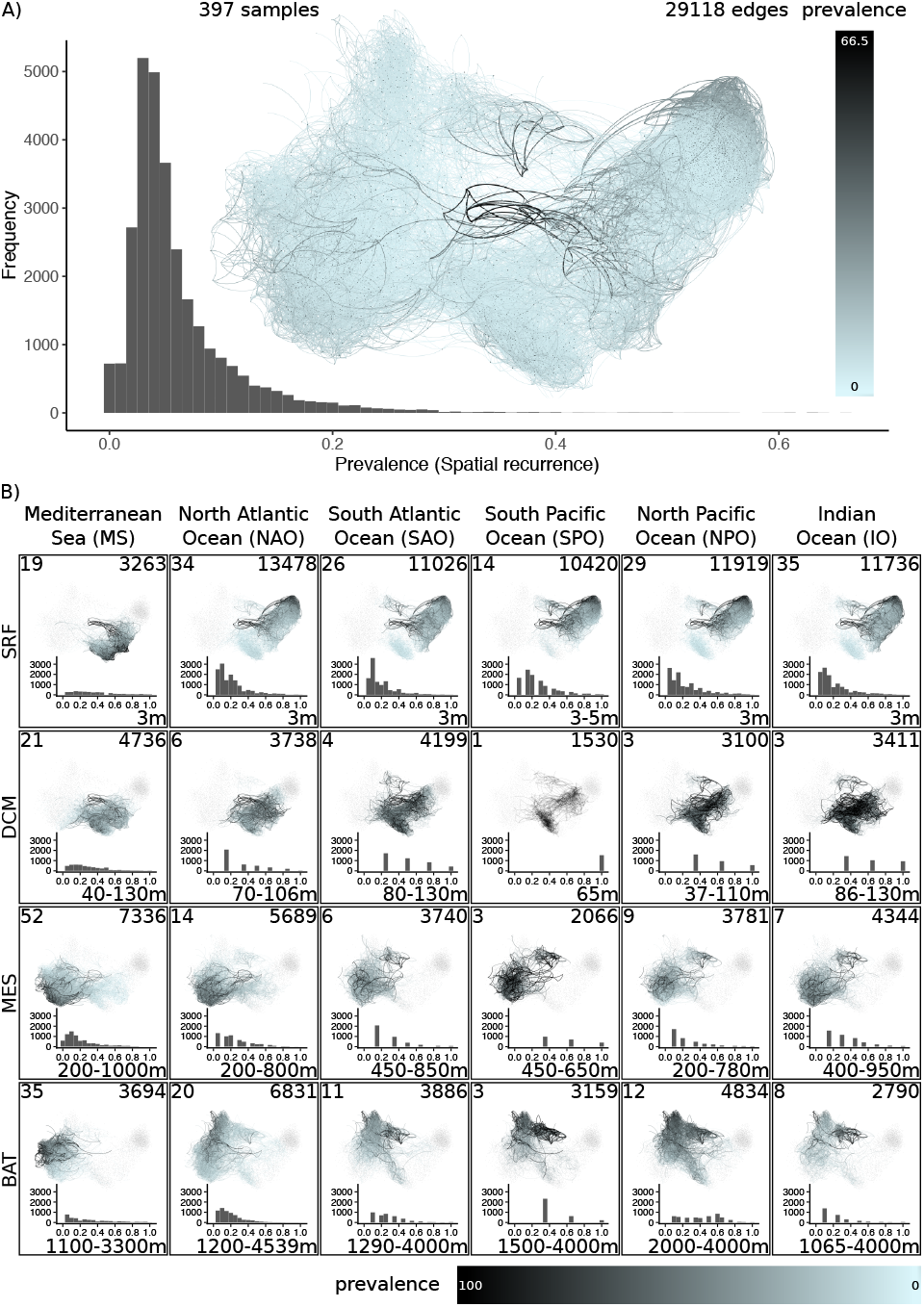
Spatial recurrence. **(A)** Association prevalence showing the fraction of subnetworks in which an association appeared considering all depth layers across the global tropical and subtropical ocean and the Mediterranean Sea. Associations that occurred more often (black) appeared in the middle of the single static network visualization. Most edges had a low prevalence (blue) <20%. **(B)** The sample-specific subnetworks of the four depth layers (rows): surface (SRF), DCM, mesopelagic (MES), and bathypelagic (BAT), in the five oceanic basins and the Mediterranean Sea (columns). The histograms show the association prevalence within each depth layer and region (excluding absent associations, i.e., 0% prevalence). The number of samples appears in the upper left corner, the number of edges with a prevalence >0% in the upper right corner, and the depth range in the lower right corner (in m below surface). Note that the prevalence goes up to 100% in **(B)** vs. 66.5% in **(A)**.

Highly prevalent associations present across all regions are candidates to represent putative core interactions in the global ocean, likely performing processes crucial for ecosystem function. We defined global associations as those appearing in more than 70% of the subnetworks in each region. In addition, we resolved prevalent (≤70% and >50%) and low-frequency (≤50% and >20%) associations. The MS is a distinct region compared with the ocean basins. For instance, the bathypelagic is warmer (median temperature of 13.8°C) than the ocean basins’ bathypelagic zone (median temperature between 1.4°C in SPO and 4.4°C in NAO). Thus, we characterized associations for all six regions, and for the ocean basins only. We found slightly to moderately more global, prevalent, and low-frequency associations when not considering the MS (Table 1, Supplementary Figure 2). The fraction of global, prevalent, and low-frequency associations was highest in the DCM layer and lowest in the bathypelagic zone (Table 1). Specifically, while we found several (28-86 without MS, and 21-26 with MS) global associations in the epi- and mesopelagic zones, only few or none (9 without MS, and none with MS) global associations were identified in the bathypelagic zone. While the epipelagic global associations were dominated by *Alphaproteobacteria*, a majority of associations from deeper zones included *Thaumarchaeota* (Supplementary Figure 2).

**Table 1:** Number of classified associations per depth layer. The sum of classified associations (including Other) is the number of present associations. Absent associations appear in other layers but in no subnetwork of a given layer. Global, prevalent, and low-frequency associations have been computed with and without considering the MS. The proportion of regional associations increased with depth (gray row).

| Depth layer | Epipelagic (Surface) | Epipelagic (DCM) | Mesopelagic | Bathypelagic |
| --- | --- | --- | --- | --- |
| Global | 26 (0.14%) | 23 (0.31%) | 21 (0.20%) | - |
| Prevalent | 22 (0.12%) | 47 (0.64%) | 10 (0.10%) | 7 (0.07%) |
| Low-frequency | 105 (0.58%) | 160 (2.17%) | 212 (2.05%) | 51 (0.51%) |
| Global (no MS) | 86 (0.47%) | 52 (0.70%) | 28 (0.27%) | 9 (0.09%) |
| Prevalent (no MS) | 207 (1.14%) | 76 (1.03%) | 27 (0.26%) | 28 (0.28%) |
| Low-frequency (no MS) | 1361 (7.46%) | 219 (2.97%) | 342 (3.30%) | 489 (4.84%) |
| Regional | 2014 (11.05%) | 2290 (31.03%) | 3420 (33.00%) | 3669 (36.33%) |
| MS | 596 (3.27%) | 1295 (17.55%) | 2254 (21.75%) | 1217 (12.05%) |
| NAO | 577 (3.16%) | 306 (4.15%) | 422 (4.07%) | 1522 (15.07%) |
| SAO | 162 (0.89%) | 304 (4.12%) | 301 (2.90%) | 143 (1.42%) |
| SPO | 152 (0.83%) | 105 (1.42%) | 40 (0.39%) | 109 (1.08%) |
| NPO | 298 (1.63%) | 133 (1.80%) | 204 (1.97%) | 516 (5.11%) |
| IO | 229 (1.26%) | 147 (1.99%) | 199 (1.92%) | 162 (1.60%) |
| Other* | 16067 (88.12%) | 4860 (65.85%) | 6701 (64.66%) | 6372 (63.10%) |
| Other (no MS)* | 14566 (79.88%) | 4743 (64.27%) | 6547 (62.17%) | 55904 (58.46%) |
| Present | 18234 (100%) | 7380 (100%) | 10364 (100%) | 10099 (100%) |
| Absent | 10884 | 21738 | 18754 | 19019 |

\*The number of unclassified (Other) associations is computed from present, regional, global, prevalent, and low-frequency associations. The last three classifications have been done with and without the MS, and subsequently the number of unclassified (other) associations varies.

### High-rank taxonomy of associations was consistent across regions

Next, we considered the most prevalent associations within a specific region and depth, i.e., those found in over 70% of the subnetworks of one region and depth layer. Despite the few global associations determined before, here, we found that high-rank taxonomic patterns of associated taxa were consistent across the water column in different regions (Figure 3). The epipelagic layers (surface and DCM) and the two lower layers (meso- and bathypelagic zones) were more similar to each other, respectively (Figure 3). The fraction of associations including *Alphaproteobacteria* was moderate to high in all zones in contrast to *Cyanobacteria* appearing mainly, as expected, in the epipelagic zone (Figure 3, Supplementary Material 1). The fraction of associations including *Dinoflagellata* was moderate to high in the epipelagic zone and lower in the meso- and bathypelagic zones (Figure 3, Supplementary Material 1). While *Dinoflagellata* associations dominated most epipelagic layers, fewer were found in the MS and SAO surface waters as well as in the DCM of the NAO (Figure 3, Supplementary Material 1). *Thaumarchaeota* associations were moderate to high especially in the mesopelagic (dominant in the MS), moderate in the bathypelagic, and low in the epipelagic zone (Figure 3, Supplementary Material 1). Associations including *Gammaproteobacteria* increased with depth, being higher in the meso- and bathypelagic than in the epipelagic, especially in the SAO, SPO, NPO and IO (Figure 3, Supplementary Material 1).

**Figure 3:**
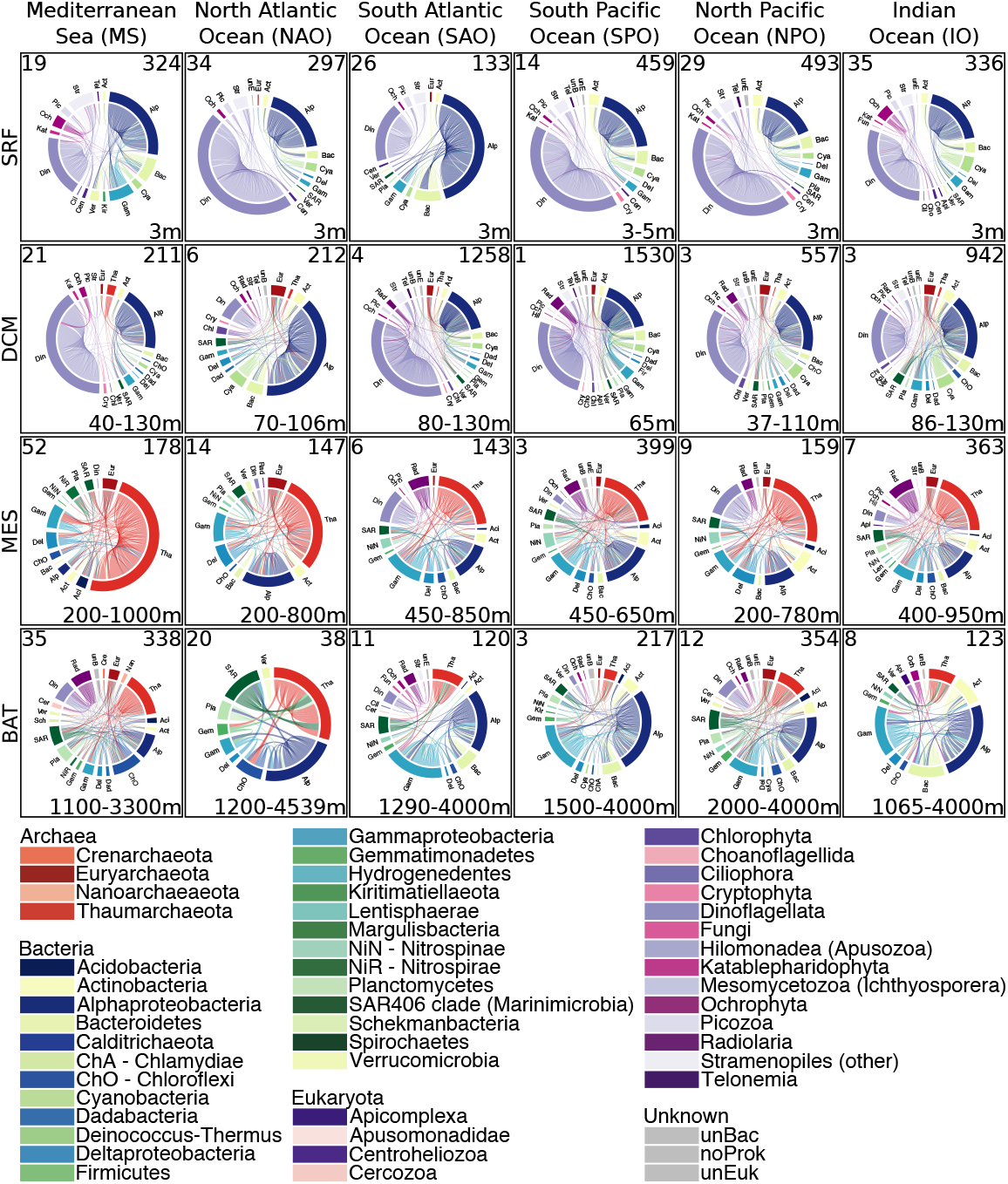
Highly prevalent associations for each region and depth layer. If an association appears in more than 70% of the subnetworks it is classified as highly prevalent. Rows indicate the four depth layers: surface (SRF), DCM, mesopelagic (MES), and bathypelagic (BAT). The number of samples appears in the upper left corner, the number of edges in the upper right corner, and the depth range in the lower right corner (in m below surface).

### The proportion of regional associations increased with depth

We determined regional associations within each depth layer. Regional associations were defined as those detected in at least one sample-specific subnetwork from one region and being absent from all subnetworks of the other five regions. Results indicated an increasing proportion of regional associations with depth (Table 1, Figure 4A-B, Supplementary Figure 3). We found substantially more associations in the DCM and mesopelagic layers of the MS than in corresponding layers of the ocean basins. This may reflect the different characteristics of these layers in the MS vs. the ocean basins or the massive differences in spatial dimensions between the ocean basins and the MS. More surface and bathypelagic regional associations were found in the MS and NAO than in other regions (Table 1). Most regional associations had low prevalence, i.e., they were present in a few sample-specific subnetworks within the region (Figure 4C). We found 235 highly prevalent (>70%) regional associations among prokaryotes, 89 among eukaryotes and 24 between domains (Supplementary Material 2).

**Figure 4:**
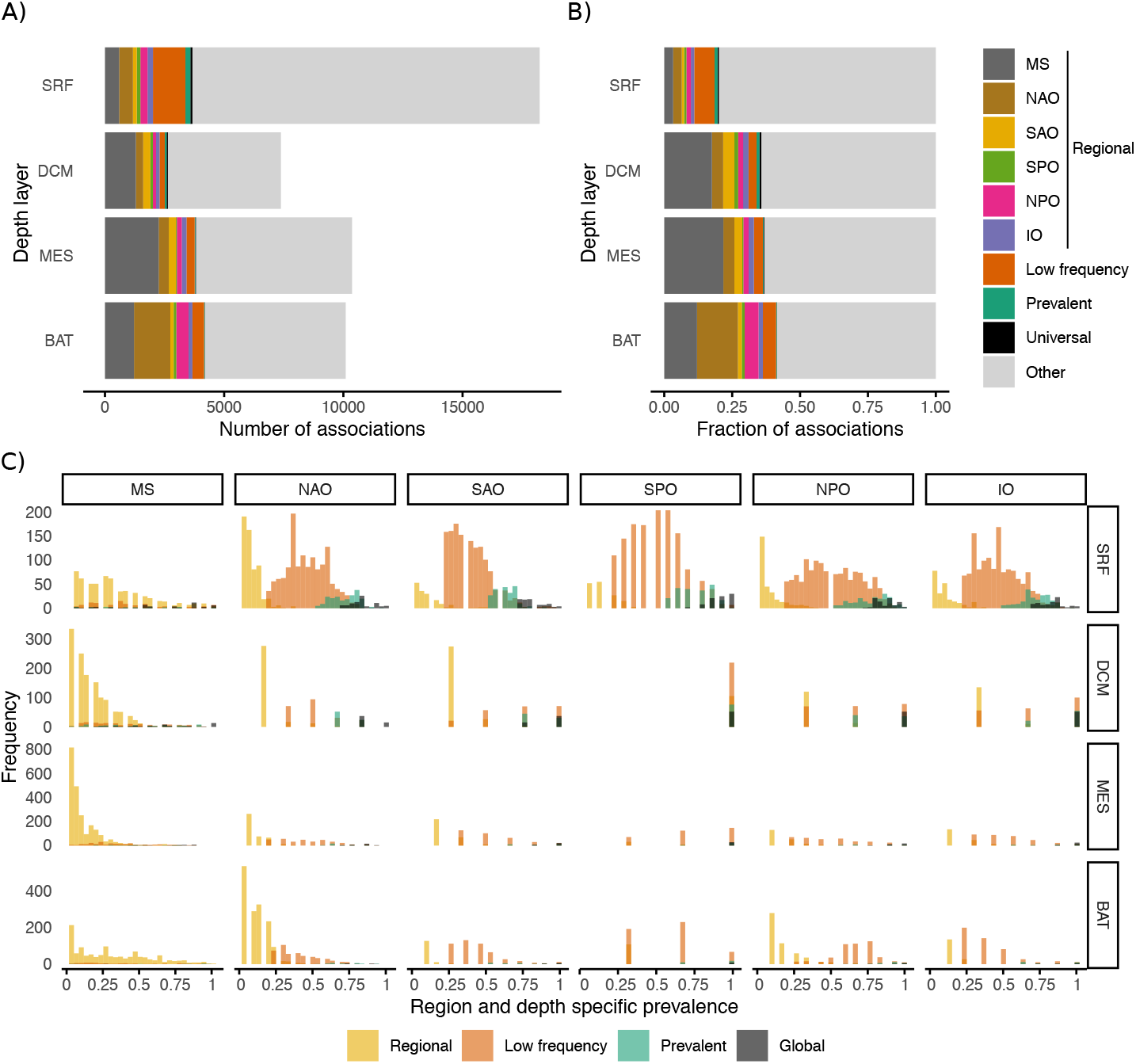
Classification of associations. We classified association into global (>70% prevalence, not considering the MS), prevalent (≤70% and >50%, not considering the MS), low-frequency (≤50% and >20%, not considering the MS), regional, and other. Regional associations are assigned to one of six ocean regions (five ocean basins and the Mediterranean Sea). The number **(A)** and fraction **(B)** of each type of association are shown for each depth layer: surface (SRF) and DCM (epipelagic), mesopelagic (MES) and bathypelagic (BAT). Color indicates the type of classification. The associations have been classified into the five types based on their prevalence in each region. The prevalence of associations is shown in **(C)**. For instance, global associations have a prevalence above 70% in each region (not considering the MS). Regional associations are present in one region (indicated with yellow with mainly low prevalence >0%) and absent in all other regions (0% prevalence not shown in graph).

### Few associations were present throughout the water column

Previous studies have found a substantial vertical connectivity in the ocean microbiota, with surface microorganisms having an impact in deep sea counterparts (11,34). Thus, here, we analyzed the vertical connectivity of potential microbial interactions, aiming to determine what surface associations could be detected along the water column. Few associations were present throughout the water column within a region, including 327 among prokaryotes, 119 among eukaryotes, and 13 between domains (Supplementary Material 3). In general, most associations from the meso- and bathypelagic did not appear in the upper layers except for the MS and NAO, where most and about half, respectively, of the bathypelagic associations already appeared in the mesopelagic (Figure 5). Specifically, 81.8 – 90.9% of the mesopelagic and 43.5-72.7% of the bathypelagic associations appeared for the first time in these layers when the five ocean basins were considered (Supplementary Table 1). In the MS, 71.2% of the mesopelagic and 22.4% of the bathypelagic associations appeared for the first time in these layers. We found that 69.7% of the associations appearing in the bathypelagic zone already appeared in the mesopelagic zone (Supplementary Table 1). This points to specific microbial interactions occurring in the deep ocean that do not occur in upper layers. In addition, most surface associations disappeared with depth in the five ocean basins and MS (Figure 5), suggesting that most surface ocean interactions are not transferred to the deep sea, despite microbial sinking (11). In fact, most deep ocean ASVs already appeared in the upper layers (Supplementary Figure 4), in agreement with previous work that has shown that a large proportion of deep sea microbial taxa are also found in surface waters, and that their presence in the deep sea is related to sinking particles (11).

**Figure 5:**
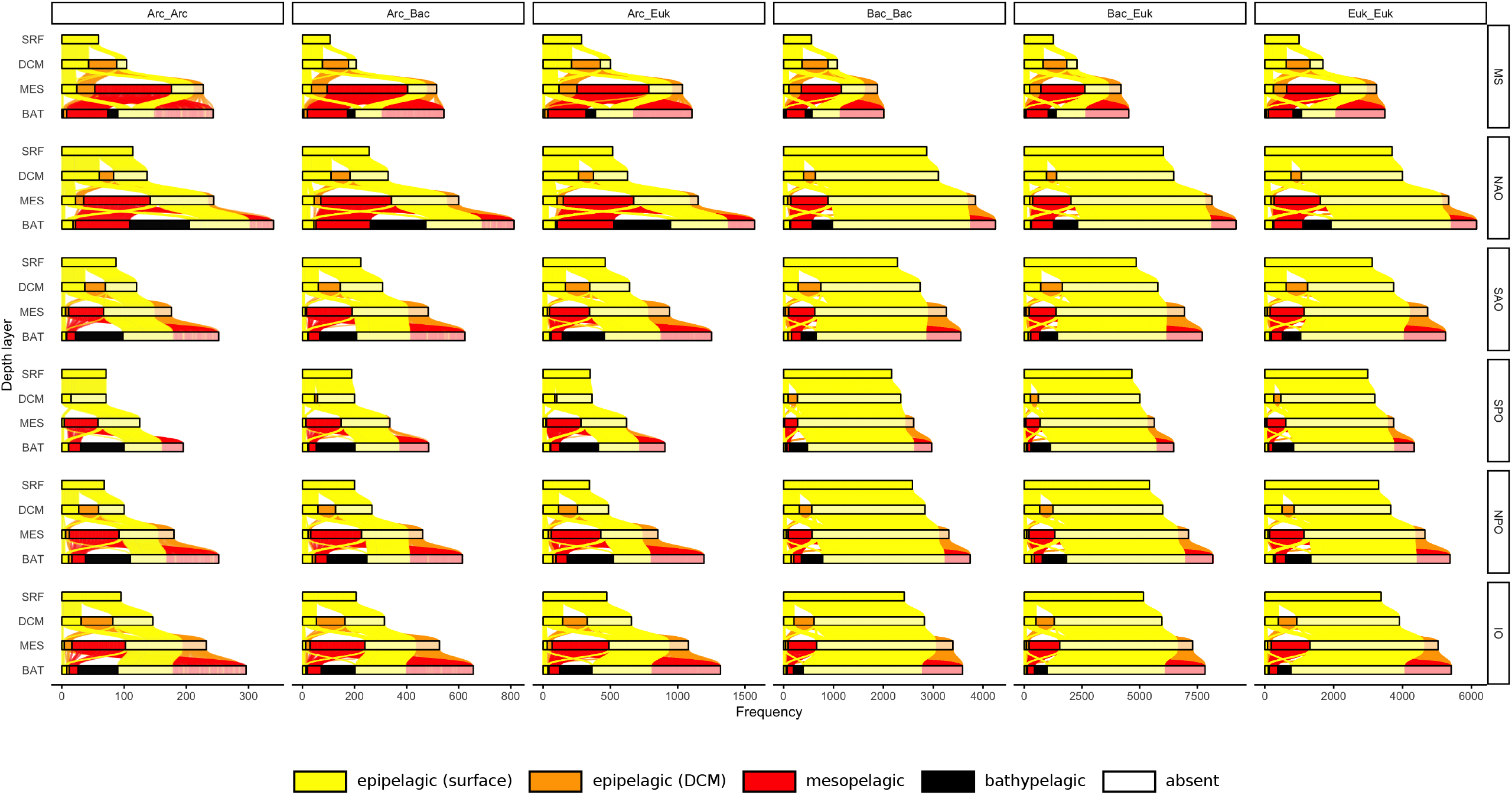
Microbial associations across depth layers. For each region and taxonomic domain, we color associations based on when they first appeared: surface (S, yellow), DCM (D, orange), mesopelagic (M, red), and bathypelagic (B, black). The SRF bar contains the associations that appeared in the surface. If they also appeared in the DCM, they are listed on the left box of the DCM bar. However, if they were not found in the DCM layer, i.e., they were absent, they appear on the right transparent box of the bar. That is, absent ASVs are grouped in the transparent box at the end of the DCM, MES, and BAT bars. Columns show associations between archaea (Arc), bacteria (Bac), and eukaryotes (Euk).

### Environmental gradients seem to shape microbial network topology

Above we grouped the sample-specific subnetworks based on regions and depth layers. However, such predefined groupings may introduce a bias to our analysis. Thus, we grouped subnetworks based on similar topology (see Methods) and identified 36 clusters of 5 to 28 subnetworks (Supplementary Table 2). We found 13 (36.1%) clusters that were dominated by surface subnetworks: six clusters (100% surface subnetworks) from three to five ocean regions but not the MS, and seven clusters including 55-86% surface networks from two to five ocean regions. In turn, 11 clusters were dominated by other layers: two DCM (64-90%), five mesopelagic (62-83%) and four bathypelagic-dominated clusters (60-69%). Nine of these 11 clusters combined different regions except for one mesopelagic and one bathypelagic-dominated cluster representing exclusively the MS (Supplementary Table 2). Furthermore, we found 11 clusters containing exclusively or mainly MS subnetworks in contrast to only one cluster dominated by an ocean basin (NAO).

Next, we built a more comprehensive representation of network similarities between subnetworks via a minimal spanning tree (MST, see Methods). The depth layers, ocean regions, location of clusters, and environmental factors were projected onto the MST (Figure 6). Most surface subnetworks were centrally located, while subnetworks from other depths appeared in different MST areas (Figure 6A). Most MS subnetworks were located in a specific branch of the MST, while the five ocean basins were mixed (Figure 6B), indicating homogeneity and connectivity within oceans but network-based differences between the oceans and the MS subnetworks. As expected, networks of the same cluster appear mostly connected in the MST (Figure 6C). Moreover, subnetworks in the MST tended to connect to subnetworks from the same depth layer or similar environmental conditions (Figure 6A, D). All in all, our results suggest a strong influence of environmental gradients, and to some extent geography, in shaping microbial network topology in the ocean (Figure 6A,B,D), as previously observed in epipelagic communities at the global scale (28).

**Figure 6:**
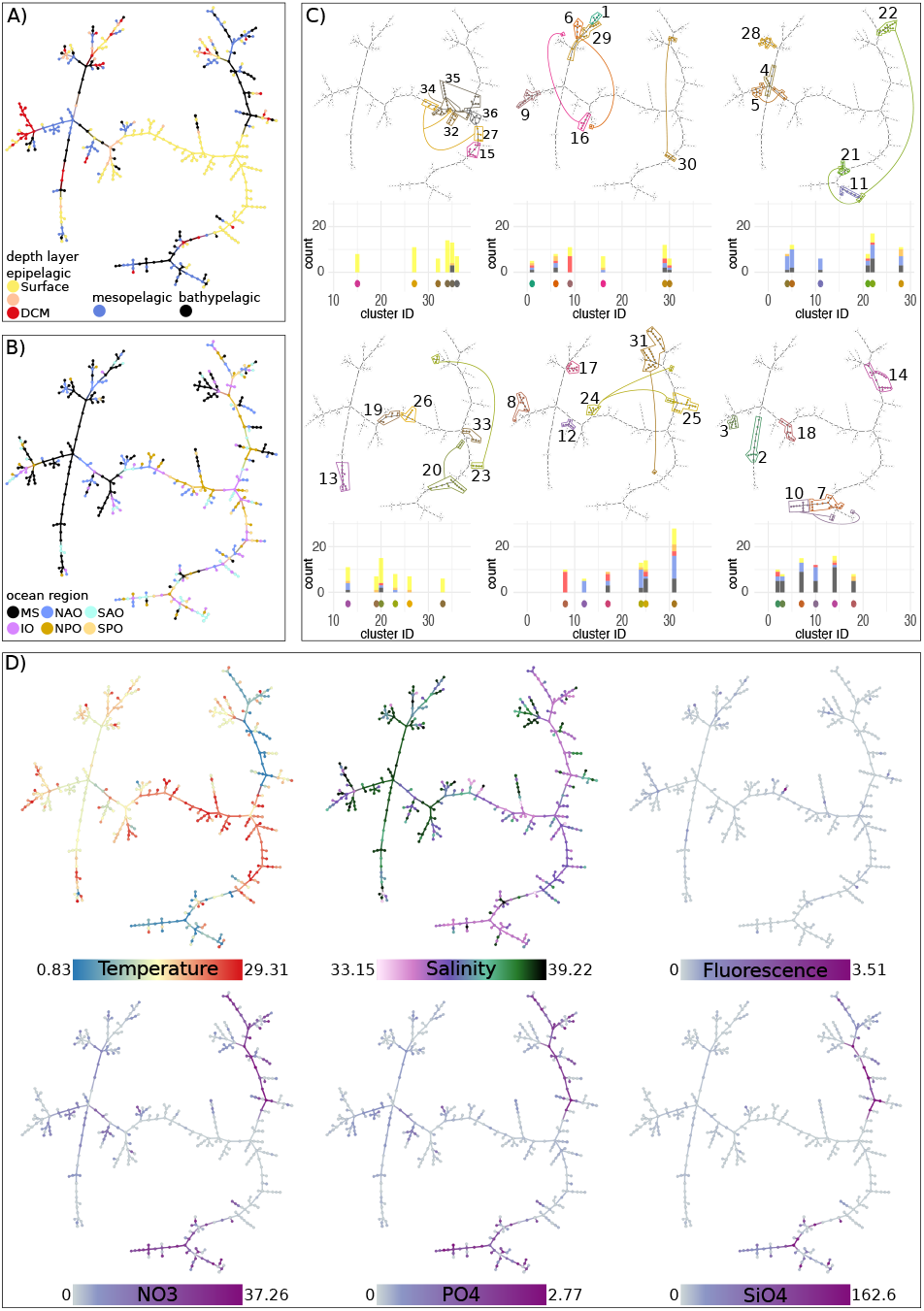
Minimal Spanning Tree. Each subnetwork is a node in the MST and represents a sample. Nodes are colored according to **(A)** the sample’s depth layer, **(B)** the sample’s ocean region, **(C)**the subnetworks cluster, and **(D)** selected environmental factors. In **(C)**, the barplots indicate the different layers within each cluster colored as in **(A)**.

## DISCUSSION

We analyzed global and regional pelagic microbial associations across the oceans’ vertical and horizontal dimensions. We found a low number of global associations indicating a potentially small global core interactome within each depth layer across the six oceanic regions. In contrast, within each region, we found less highly prevalent associations in the bathypelagic zone of the global ocean (pointing to a smaller regional core) than in the upper layers, except from the NPO, which had less highly prevalent associations in the meso-than in the bathypelagic. In turn, we found more regional associations in the bathypelagic than in upper layers. This may reflect the heterogeneity and isolation of deep ocean regions due to deep currents, water masses, or the topography of the seafloor that may prevent microbial dispersal. Moreover, the higher complexity of the deep ocean ecosystem may provide a higher number of ecological niches potentially resulting in more regional associations. Niche diversification may be associated to the quality and types (labile, recalcitrant, etc.) of organic matter reaching the deep ocean from the epipelagic zone (29), which is significantly different across oceanic regions (35). In an exploration of generalists versus specialist prokaryotic metagenome-assembled genomes (MAGs) in the arctic Ocean, most of the specialists were linked to mesopelagic samples indicating that their distribution was uneven across depth layers (36). This is in agreement with putatively more niches in the deep ocean than in upper ocean layers leading to more specialist taxa and subsequently more regional associations in deep ocean waters.

Vertical connectivity in the ocean microbiome is partially modulated by surface productivity through sinking particles (11,34,37). An analysis of eight stations, distributed across the Atlantic, Pacific and Indian oceans (including 4 depths: Surface, DCM, meso- and bathypelagic), indicated that bathypelagic communities comprise both endemic taxa as well as surface-related taxa arriving via sinking particles (11). Another work (34) identified for both components (i.e. surface-related and deep-endemic) the dominating phylogenetic groups: while *Thaumarchaeota, Deltaproteobacteria, OM190* (*Planctomycetes*) and *Planctomycetacia* (*Planctomycetes*) dominated the endemic bathypelagic communities, *Actinobacteria, Alphaproteobacteria, Gammaproteobacteria* and *Flavobacteriia* (*Bacteroidetes*) dominated the surface-related taxa in the bathypelagic zone. We found association partners for each dominating phylogenetic group within each investigated type of association, i.e., highly prevalent, regional, global, prevalent, and low-frequency associations. While ASVs belonging to these taxonomic groups were present throughout the water column, specific associations were observed especially in the mesopelagic and bathypelagic zones, which suggests specific interactions between endemic deep-sea taxa, in agreement with the hypothesis indicating high niche partitioning and more specialist taxa in the deep ocean (38,39). This is in agreement with a recent study that found a remarkable taxonomic novelty in the deep ocean after analyzing 58 microbial metagenomes from a global deep-sea survey, unveiling ∼68% archaeal and ∼58% bacterial novel species (40).

Little is known about the distribution of microbial interactions across the water column. Associations found along the entire water column could point to microbes interacting across all water layers or interacting microbes that sink together (41). We found that associations present across all layers were limited, pointing to a heterogeneous distribution of interactions in the water column. Given that we targeted the picoplankton, the associated taxa found in the entire water column may represent non-physical interactions occurring in all water layers, instead of interactions occurring in sinking particles (41). A fraction of the associations observed only in the deep ocean may correspond to microbial consortia degrading sinking particles, or taxa that might have detached from sinking particles, i.e., dual life-style taxa as observed in (42). Altogether, our results suggest that most microbial interactions change across the water column, while a few are maintained. Furthermore, some microorganisms may change their interaction partners across the water column. Changes of microbial interactions with depth could also be linked to ecological successions in sinking particles (43), yet our spatial sampling precludes us from investigating this possibility.

In our study, mesopelagic subnetworks displayed the lowest network connectivity (determined via edge density) across most regions on average, and we found the strongest associations among both meso- and bathypelagic subnetworks. Moreover, we found the highest clustering (transitivity) in the meso- and bathypelagic zones (relatively colder waters) compared to the epipelagic zone (warmer waters). Similarly, a previous global-scale study (28) concentrating on the epipelagic zone and including polar waters, found higher edge density, association strength and clustering in polar waters compared to warmer waters. These results suggest that either microorganisms interact more in colder environments or that their recurrence is higher due to a higher environmental selection exerted by low temperatures. Alternatively, limited resources (primarily nutrients) in the surface versus the deep tropical and subtropical ocean may prevent the establishment of specific microbial interactions in surface waters. Furthermore, environmental stability in the deep sea may have led to high niche partitioning (38,39), which could have promoted the establishment of interactions in the meso- and bathypelagic.

Through quantifying regional associations, our results indicated distinct associations in the MS, where most regional associations were observed compared to the ocean basins, as previously shown in an epipelagic network (26). The Mediterranean Sea is a hotspot of multicellular biodiversity and endemic species (44,45), and despite being less studied than animals and plants, there are also reports of putatively endemic microorganisms, such as specific SAR11 (46). Thus, part of the recovered associations could be reflecting endemic interactions derived from endemic as well as non-endemic taxa. Potentially endemic taxa should be investigated at the genome level, given that the 16S or 18S may not reflect fine-grained differences (47,48). Furthermore, we found a substantial number of regional associations in the NAO compared to other ocean basins, contrasting with the NAO having the lowest number of regional associations in a previous epipelagic network (26). Given that the previous studies used different samples, these results are not surprising.

To conclude, we have disentangled the spatial distribution of associations in the global ocean microbiome, from surface to bottom water layers, finding both global and regional microbial associations. Our analysis captured network topology changes across vertical (water column) and horizontal (different regions) pelagic zones of the ocean. Furthermore, our results indicate that associations have specific biogeographies that do not necessarily mirror taxonomic biogeographies.

## METHODS

### Dataset

Samples originated from two expeditions, Malaspina-2010 (49) and Hotmix (50). The former was onboard the R/V Hespérides and most ocean basins were sampled between December 2010 and July 2011. Malaspina samples included i) *MalaSurf*, surface samples (5,51), ii) *MalaVP*, vertical profiles (14), and iii) *MalaDeep*, deep-sea samples, (52–54). In the Hotmix expedition, sampling took place onboard the R/V Sarmiento de Gamboa between 27th April and 29th May 2014 and represented a quasi-synoptic transect across the MS and the adjacent North-East of the NAO. See details in Table 2.

**Table 2:** Used datasets. We required that each sample had to provide data for both eukaryotes and prokaryotes, which resulted in 397 samples. This condition allowed only 13 MalaDeep samples. 16S and 18S refer to sequenced samples.

| Dataset | Samples used for analysis | Stations | Depth range (m) | Water samples | Size Fraction (µm) | 16S | 18S | Reference | ENA accession number |
| --- | --- | --- | --- | --- | --- | --- | --- | --- | --- |
| *Malaspina* |  |  |  |  |  |  |  | (5,51) |  |
| *MalaSurf* | 122 | 120 | 3 | 122 | 0.2-3 | 122 | 124 | (5,51) | PRJEB23913 [18S rRNA genes],<br>PRJEB25224 [16S rRNA genes] |
| *MalaVP* | 83 | 13 | 3-4000 | 91 | 0.2-3 | 91 | 83 | (14) & This study | PRJEB23771 [18S rRNA genes],<br>PRJEB45015 [16S rRNA genes] |
| *MalaDeep (Prok\*)* | 13 | 30 | ~4000 | 60 | 0.2-0.8 | 41 | - | (54) | PRJEB45011 |
| *MalaDeep (Euk\*)* | 13 | 27 | 2400-4000 | 27 | 0.8-20 | - | 82 | This study | PRJEB45014 |
| Hotmix | 179 | 29 | 3-4539 | 188 | 0.2-3 | 188 | 179 | (60) | PRJEB44683 [18S rRNA genes],<br>PRJEB44474 [16S rRNA genes] |

\*Prok - prokaryotes; Euk - eukaryotes

DNA extractions are indicated in the publications associated with each dataset (Table 2). The 16S and 18S rRNA genes were amplified and sequenced. PCR amplification and sequencing of *MalaSurf, MalaVP* (18S), and *Hotmix* (16S) are indicated in the publications associated with each dataset in Table 2. *MalaVP* (16S) and *Hotmix* (18S) were PCR-amplified and sequenced following the same approach as in (5). The DNA from *MalaDeep* samples was extracted as indicated in (52,53) and re-sequenced at Genoscope (France) with the primers indicated below. *MalaSurf, MalaVP* and *Hotmix* datasets were sequenced at RTL Genomics (Texas, USA).

We used the same amplification primers for all samples. For the 16S, we amplified the V4-V5 hypervariable region using the primers 515F-Y and 926R (55). For the 18S, we amplified the V4 hypervariable region with the primers TAReukFWD1 and TAReukREV3 (56). See more details in (5). Amplicons were sequenced in *Illumina* MiSeq or HiSeq2500 platforms (2×250 or 2×300 bp reads). Operational Taxonomic Units were delineated as Amplicon Sequence Variants (ASVs) using DADA2 (57), running each dataset separately before merging the results. ASVs were assigned taxonomy using SILVA (58), v132, for prokaryotes, and PR2 (59), v4.11.1, for eukaryotes. ASVs corresponding to Plastids, Mitochondria, Metazoa, and Plantae, were removed. Only samples with at least 2000 reads were kept. The dataset contained *MalaDeep* replicates, which were merged, and two filter size fractions: given the cell sizes of prokaryotes versus microeukaryotes, we used the smallest size-fraction (0.2-0.8 µm) for prokaryotes and the larger one (0.8-20 µm) for microbial eukaryotes. The other three datasets considered the 0.2-3 µm size fraction. Additionally, we required that samples had eukaryotic and prokaryotic data, resulting in 397 samples for downstream analysis: 122 *MalaSurf*, 83 *MalaVP*, 13 *MalaDeep*, and 179 *Hotmix* (Table 2). We separated the samples into epipelagic, mesopelagic and bathypelagic zone (Figure 1). Furthermore, we separated most epipelagic zone samples into surface layer and deep-chlorophyll maximum (DCM) layer, but 18 MS and 4 NAO samples belonged to neither. We also considered environmental variables: Temperature (2 missing values = mv), salinity (2 mv), fluorescence (3 mv), and inorganic nutrients NO_3_^−^ (36 mv), PO_4_^3−^ (38 mv), and SiO_2_ (37 mv), which were measured as indicated elsewhere (5,14,60). In specific samples, missing data on nutrient concentrations were estimated from the World Ocean Database (61).

### Single static network

We constructed the single static network in four steps. First, we prepared the data for network construction. We excluded rare microorganisms by keeping ASVs with a sequence abundance sum above 100 reads across all samples and appearing in at least 20 samples (>5% of the dataset). The latter condition removed larger eukaryotes only appearing in the 13 *MalaDeep* eukaryotic samples of the 0.8-20 µm size fraction. To control for data compositionality (62), we applied a centered-log-ratio transformation separately to the prokaryotic and eukaryotic tables before merging them.

Second, we inferred a (preliminary) network using FlashWeave (63), selecting the options “heterogeneous” and “sensitive”. FlashWeave was chosen as it can handle sparse datasets like ours, taking zeros into account and avoiding spurious correlations between ASVs that share many zeros. This initial network had 5457 nodes and 31,966 edges, 30,657 (95.9%) positive and 1309 (4.1%) negative.

Third, we aimed at removing environmentally-driven edges. FlashWeave can detect indirect edges and can also consider metadata such as environmental variables, but currently does not support missing data. Thus, we applied EnDED (64), combining the methods Interaction Information (with 0.05 significance threshold and 10,000 iterations) and Data Processing Inequality as done previously via artificially-inserted edges to connect all microbial nodes to the six environmental parameters (33). Although EnDED can handle missing environmental data when calculating intermediate values relating ASV and environmental factors, it would compute intermediate values for microbial edges using all samples. Thus, to avoid a possible bias and speed up the calculation process, we applied EnDED individually for each environmental factor, using only the samples containing values for the specific environmental factor. We detected and removed potential environmentally-driven edges due to nutrients (4.9% NO_3_^−^, 4.2% PO_4_^3−^, 2.0% SiO_2_), temperature (1.9%), salinity (0.2%), and Fluorescence (0.01%) (Supplementary Table 3).

Fourth, we removed isolated nodes, i.e., nodes without any edge. The resulting network represented the single static network in our study. It contained 5448 nodes and 29,118 edges; 28,178 (96.8%) positive and 940 (3.2%) negative.

### Sample-specific subnetwork

We constructed 397 sample-specific subnetworks. Each subnetwork represented one sample and was derived from the single static network, i.e., a subnetwork contained nodes and edges present in the single static network but not vice versa. First, we required that an edge must be present in the single static network. Second, an edge can only be present within a subnetwork if both microorganisms associated with the edge have a sequence abundance above zero in the corresponding sample. Third, microorganisms associated need to appear together (intersection) in more than 20% of the samples, in which one or both appear (union) for a specific region and depth.

Formally, consider sample *s*_*RL*_ with *R* being the marine region, and *L* the sample’s depth layer. Let *e* be an association between microorganisms *A* and *B*. Then, association *e* is present in the sample-specific subnetwork *N*_*s*_, if

i. *e* is an association in the single static network,
ii. the microorganisms *A* and *B* are present within sample *s*, i.e., the abundances are above zero within that particular sample, and
iii. the association has a region and depth specific Jaccard index, *J*_*RL*_, above 20% (see below).

In addition to these three conditions, a node is present in a sample-specific subnetwork when connected to at least one edge, i.e., we removed isolated nodes.

Regarding the third condition, we determined *J*_*RL*_ for each association pair by computing within each region and depth layer, the fraction of samples two microorganisms appeared together (intersection) from the total samples at least one microorganism appears (union). Supplementary Table 4 shows the number of edges using different thresholds. Given the heterogeneity of the dataset within regions and depth layers, we decided to use a low threshold, keeping edges with a Jaccard index above 20% and removed edges below or equal to 20%. The third condition was robust (Supplementary Figure 5). We tested robustness by randomly drawing a subset of samples from each region and depth combination. The subset contained between 10% and 90% of the original samples. We rounded up decimal numbers to avoid zero sample subsets, e.g., 10% of 7 samples results in a subset of 1 sample. We excluded the DCM of the SPO because it contained only one sample. Next, we recomputed the Jaccard index for the random subset. Lastly, requiring J>20%, we evaluated robustness determining i) how many edges were kept in the random subsamples compared to all samples, and ii) how many edges were kept in the random subset that were also kept when all samples were used. We repeated the procedure for each region-depth combination 1000 times.

### Spatial recurrence

To determine an association’s spatial recurrence, we calculated its prevalence as the fraction of subnetworks in which the association was present. We determined association prevalence across the 397 samples and each region-layer combination. We mapped the scores onto the single static network, visualized in Gephi (65) v.0.9.2, using the Fruchterman Reingold Layout (66) with a low gravity score of 0.5. We used the region-layer prevalence to determine global and regional associations. We considered an association to be global within a specific depth layer if its prevalence was above 70% in all regions. In turn, a regional association had an association prevalence above 0% within a particular region-layer (present, appearing in at least one subnetwork) and 0% within other regions of the same layer (absent, appearing in no subnetwork). We further characterized associations that were neither global nor local. We considered an association to be prevalent within a specific depth layer if its prevalence was above 50% in all regions. Similarly, associations that appear in a specific depth layer in all regions over 20% are considered low-frequency. Thus, an association can be classified as i) global, ii) regional, iii) prevalent, iv) low-frequency, and v) “other”, i.e., associations that have not been classified into the previous categories.

### Network metrics

We considered the *number of nodes* and *edges* and six other network metrics of which most were computed with functions of the igraph R-package (67). *Edge density* indicating connectivity is computed through the number of actual edges divided by the number of possible edges. The *average path length* is the average length of all shortest paths between nodes in a network. *Transitivity*, indicating how well a network is clustered, is the probability that the nodes’ neighbors are connected. *Assortativity* measures if similar nodes tend to be connected, i.e., *assortativity (degree)* is positive if high degree nodes tend to connect to other high degree nodes and negative otherwise.

Similarly, *assortativity (Euk-Prok)* is positive if eukaryotes tend to connect to other eukaryotes while prokaryotes tend to connect to other prokaryotes. Lastly, we computed the *average positive association strength* as the mean of all positive association scores provided by Flash Weave.

### Similar networks based on network topology

The previous metrics (so-called global network metrics) disregard local structures’ complexity, and topological analyses should include local metrics (68), e.g., graphlets (69). Here, we determined network-dissimilarity between each pair of sample-specific subnetworks as proposed in (70), comparing network topology without considering specific ASVs. The network-dissimilarity is a distance measurement that is always positive: 0 if networks are identical and greater numbers indicate greater dissimilarity.

Next, we constructed a Network Similarity Network (NSN), where each node is a subnetwork and each node connects with all other nodes, i.e., the NSN was a complete graph. We assigned the network-dissimilarity score as edge weight within the NSN. To simplify the NSN while preserving its main patterns, we determined the minimal spanning tree (MST) of the NSN. The MST had 397 nodes and 396 edges. The MST is a backbone, with no circular path, in which the edges are chosen so that the edge weights sum is minimal and all nodes are connected, i.e., a path exists between any two nodes. We determined the MST using the function *mst* in the igraph package in R (67,71).

Using the network-dissimilarity (distance) matrix, we determined clusters of similar subnetworks using Python scripts. First, we reduced the matrix to ten dimension using *umap* (72) with the following parameter settings: n_neighbors=3, min_dist=0, n_components=10, random_state=123, and metric=’precomputed’. Second, we clustered the subnetworks (represented via ten dimensions) with *hdbscan* (73) setting the parameters to min_samples=3 and min_clusters=5.

## Acknowledgements

We thank all members of the Malaspina and Hotmix expeditions and the multiple projects funding these collaborative efforts. Sampling was carried out thanks to the Consolider-Ingenio programme (project Malaspina 2010 Expedition, ref. CSD2008–00077) and HOTMIX project (CTM2011-30010/MAR), Competitiveness Science and Innovation. Part of the analyses have been performed at the Marbits bioinformatics core at ICM-CSIC (https://marbits.icm.csic.es). This project and IMD received funding from the European Union′s Horizon 2020 research and innovation program under the Marie Skłodowska-Curie grant agreement no. 675752 (ESR2, http://www.singek.eu) to RL. RL was supported by a Ramón y Cajal fellowship (RYC-2013-12554, MINECO, Spain). This work was also supported by the projects INTERACTOMICS (CTM2015-69936-P, MINECO, Spain), MicroEcoSystems (240904, RCN, Norway) and MINIME (PID2019-105775RB-I00, AEI, Spain) to RL. SC was supported by the CNRS MITI through the interdisciplinary program Modélisation du Vivant (GOBITMAP grant). SC, DE and SGA were funded by the H2020 project AtlantECO (award number 862923). We acknowledge funding of the Spanish government through the ‘Severo Ochoa Centre of Excellence’ accreditation (CEX2019-000928-S).

## Author’s contributions

The overall project was conceived and designed by RL. JMG, CMD, SGA, RM, JA were responsible for the sampling and acquisition of contextual data. CRG, JP and MS processed specific samples in the laboratory. RL processed the amplicon data generating the two ASV tables. They were the starting point of the present study, which is part of the overall project. IMD developed the conceptual approach and DE, SC, and RL contributed to its finalization. IMD performed the data analysis. ED, MS, CMD, SGA, RM, JMG, DE, SC, and RL contributed with interpretation of the results. IMD wrote the original draft. All authors contributed to manuscript revisions and approved the final version of the manuscript.

## Competing interests

The authors declare that they have no competing interests.

## Data availability and Reproducibility

Sequence data is publicly available at the European Nucleotide Archive (see accession numbers in Table 2). The code for data analysis including commands to run FlashWeave and EnDED (environmentally-driven-edge-detection and computing Jaccard index) are publicly available: https://github.com/InaMariaDeutschmann/GlobalNetworkMalaspinaHotmix.

## Notes

### Competing Interest Statement

The authors have declared no competing interest.

### Summary of Updates

We revised the text especially discussion, and revised figures (now 6 main and 5 supplementary figures)

## REFERENCES

1. Falkowski PG, Fenchel T, Delong EF. The Microbial Engines That Drive Earth’s Biogeochemical Cycles. Vol. 320, Science. American Association for the Advancement of Science; 2008. p. 1034–9.

2. DeLong EF. The microbial ocean from genomes to biomes. Vol. 459, Nature. 2009. p. 200–6.

3. Krabberød AK, Bjorbækmo MFM, Shalchian-Tabrizi K, Logares R. Exploring the oceanic microeukaryotic interactome with metaomics approaches. Aquatic Microbial Ecology. 2017;79(1):1–12.

4. Vellend M. The theory of ecological communities (MPB-57). Princeton University Press; 2020.

5. Logares R, Deutschmann IM, Junger PC, Giner CR, Krabberød AK, Schmidt TSB, et al. Disentangling the mechanisms shaping the surface ocean microbiota. Microbiome. 2020;8(1):55.

6. Sunagawa S, Coelho LP, Chaffron S, Kultima JR, Labadie K, Salazar G, et al. Structure and function of the global ocean microbiome. Science. 2015 May 22;348(6237):1261359.

7. Ibarbalz FM, Henry N, Brandão MC, Martini S, Busseni G, Byrne H, et al. Global Trends in Marine Plankton Diversity across Kingdoms of Life. Cell. 2019;179(5):1084-1097.e21.

8. Salazar G, Paoli L, Alberti A, Huerta-Cepas J, Ruscheweyh HJ, Cuenca M, et al. Gene Expression Changes and Community Turnover Differentially Shape the Global Ocean Metatranscriptome. Cell. 2019;179(5):1068-1083.e21.

9. Cram JA, Xia LC, Needham DM, Sachdeva R, Sun F, Fuhrman JA. Cross-depth analysis of marine bacterial networks suggests downward propagation of temporal changes. The ISME Journal. 2015;9(12):2573–86.

10. Parada AE, Fuhrman JA. Marine archaeal dynamics and interactions with the microbial community over 5 years from surface to seafloor. The ISME Journal. 2017;11(11):2510–25.

11. Mestre M, Ruiz-González C, Logares R, Duarte CM, Gasol JM, Sala MM. Sinking particles promote vertical connectivity in the ocean microbiome. Proc Natl Acad Sci USA. 2018 Jul 17;115(29):E6799.

12. Peoples LM, Donaldson S, Osuntokun O, Xia Q, Nelson A, Blanton J, et al. Vertically distinct microbial communities in the Mariana and Kermadec trenches. PLOS ONE. 2018;13(4):1–21.

13. Xu Z, Wang M, Wu W, Li Y, Liu Q, Han Y, et al. Vertical Distribution of Microbial Eukaryotes From Surface to the Hadal Zone of the Mariana Trench. Frontiers in Microbiology. 2018;9:2023.

14. Giner CR, Pernice MC, Balagué V, Duarte CM, Gasol JM, Logares R, et al. Marked changes in diversity and relative activity of picoeukaryotes with depth in the world ocean. The ISME Journal. 2020 Feb 1;14(2):437–49.

15. Massana R, Logares R. Eukaryotic versus prokaryotic marine picoplankton ecology. Environmental Microbiology. 2013;15(5):1254–61.

16. Layeghifard M, Hwang DM, Guttman DS. Disentangling Interactions in the Microbiome: A Network Perspective. Vol. 25, Trends in Microbiology. 2017. p. 217–28.

17. Seymour JR, Amin SA, Raina JB, Stocker R. Zooming in on the phycosphere: the ecological interface for phytoplankton–bacteria relationships. Nature Microbiology. 2017;2(7):17065.

18. Bjorbækmo MFM, Evenstad A, Røsæg LL, Krabberød AK, Logares R. The planktonic protist interactome: where do we stand after a century of research? The ISME Journal [Internet]. 2019; Available from: https://doi.org/10.1038/s41396-019-0542-5

19. Baldauf SL. An overview of the phylogeny and diversity of eukaryotes. Journal of Systematics and Evolution. 2008;46(3):263.

20. Lewis WH, Tahon G, Geesink P, Sousa DZ, Ettema TJG. Innovations to culturing the uncultured microbial majority. Nature Reviews Microbiology [Internet]. 2020 Oct 22; Available from: https://doi.org/10.1038/s41579-020-00458-8

21. Steele JA, Countway PD, Xia L, Vigil PD, Beman JM, Kim DY, et al. Marine bacterial, archaeal and protistan association networks reveal ecological linkages. The ISME Journal. 2011;5(9):1414–25.

22. Chow CET, Sachdeva R, Cram JA, Steele JA, Needham DM, Patel A, et al. Temporal variability and coherence of euphotic zone bacterial communities over a decade in the Southern California Bight. The ISME Journal. 2013;7(12):2259–73.

23. Chow CET, Kim DY, Sachdeva R, Caron DA, Fuhrman JA. Top-down controls on bacterial community structure: microbial network analysis of bacteria, T4-like viruses and protists. The ISME Journal. 2014;8(4):816–29.

24. Needham DM, Sachdeva R, Fuhrman JA. Ecological dynamics and co-occurrence among marine phytoplankton, bacteria and myoviruses shows microdiversity matters. The ISME Journal. 2017;11(7):1614– 29.

25. Krabberød AK, Deutschmann IM, Bjorbækmo MFM, Balagué V, Giner CR, Ferrera I, et al. Long-term patterns of an interconnected core marine microbiota. Environmental Microbiome. 2022 May 7;17(1):22.

26. Lima-Mendez G, Faust K, Henry N, Decelle J, Colin S, Carcillo F, et al. Determinants of community structure in the global plankton interactome. Science. 2015;348(6237):1262073.

27. Milici M, Deng ZL, Tomasch J, Decelle J, Wos-Oxley ML, Wang H, et al. Co-occurrence Analysis of Microbial Taxa in the Atlantic Ocean Reveals High Connectivity in the Free-Living Bacterioplankton. Frontiers in Microbiology. 2016;7:649.

28. Chaffron S, Delage E, Budinich M, Vintache D, Henry N, Nef C, et al. Environmental vulnerability of the global ocean epipelagic plankton community interactome. Sci Adv. 2021 Aug;7(35).

29. Arístegui J, Gasol JM, Duarte CM, Herndld GJ. Microbial oceanography of the dark ocean’s pelagic realm. Limnology and Oceanography. 2009;54(5):1501–29.

30. Shade A, Handelsman J. Beyond the Venn diagram: the hunt for a core microbiome. Environmental Microbiology. 2012;14(1):4–12.

31. Coutinho FH, Meirelles PM, Moreira APB, Paranhos RP, Dutilh BE, Thompson FL. Niche distribution and influence of environmental parameters in marine microbial communities: a systematic review. PeerJ. 2015 Jun;3:e1008.

32. Mandakovic D, Rojas C, Maldonado J, Latorre M, Travisany D, Delage E, et al. Structure and co-occurrence patterns in microbial communities under acute environmental stress reveal ecological factors fostering resilience. Scientific Reports. 2018;8(1):5875.

33. Deutschmann IM, Krabberød AK, Latorre F, Delage E, Marrasé C, Balagué V, et al. Disentangling temporal associations in marine microbial networks. bioRxiv. 2022 Jan 1;2021.07.13.452187.

34. Ruiz-González C, Mestre M, Estrada M, Sebastián M, Salazar G, Agustí S, et al. Major imprint of surface plankton on deep ocean prokaryotic structure and activity. Molecular Ecology. 2020;29(10):1820–38.

35. Hansell DA, Carlson CA. Deep-ocean gradients in the concentration of dissolved organic carbon. Nature. 1998 Sep 1;395(6699):263–6.

36. Royo-Llonch M, Sánchez P, Ruiz-González C, Salazar G, Pedrós-Alió C, Sebastián M, et al. Compendium of 530 metagenome-assembled bacterial and archaeal genomes from the polar Arctic Ocean. Nature Microbiology. 2021 Dec 1;6(12):1561–74.

37. Boeuf D, Edwards BR, Eppley JM, Hu SK, Poff KE, Romano AE, et al. Biological composition and microbial dynamics of sinking particulate organic matter at abyssal depths in the oligotrophic open ocean. Proc Natl Acad Sci USA. 2019 Jun 11;116(24):11824.

38. McClain CR, Schlacher TA. On some hypotheses of diversity of animal life at great depths on the sea floor. Marine Ecology. 2015 Dec 1;36(4):849–72.

39. R. Hessler R, L. Sanders H. Faunal diversity in the deep-sea. Deep Sea Research and Oceanographic Abstracts. 1967 Feb 1;14(1):65–78.

40. Acinas SG, Sánchez P, Salazar G, Cornejo-Castillo FM, Sebastián M, Logares R, et al. Deep ocean metagenomes provide insight into the metabolic architecture of bathypelagic microbial communities. Communications Biology. 2021 May 21;4(1):604.

41. Bochdansky AB, Clouse MA, Herndl GJ. Eukaryotic microbes, principally fungi and labyrinthulomycetes, dominate biomass on bathypelagic marine snow. ISME J. 2017 Feb;11(2):362–73.

42. Sebastián M, Sánchez P, Salazar G, Álvarez-Salgado XA, Reche I, Morán XAG, et al. The quality of dissolved organic matter shapes the biogeography of the active bathypelagic microbiome. bioRxiv. 2021 Jan 1;2021.05.14.444136.

43. Pelve EA, Fontanez KM, DeLong EF. Bacterial Succession on Sinking Particles in the Ocean’s Interior. Frontiers in Microbiology [Internet]. 2017;8. Available from: https://www.frontiersin.org/articles/10.3389/fmicb.2017.02269

44. Coll M, Piroddi C, Steenbeek J, Kaschner K, Ben Rais Lasram F, Aguzzi J, et al. The biodiversity of the Mediterranean Sea: estimates, patterns, and threats. PLoS One. 2010 Aug 2;5(8):e11842.

45. Danovaro R, Company JB, Corinaldesi C, D’Onghia G, Galil B, Gambi C, et al. Deep-sea biodiversity in the Mediterranean Sea: the known, the unknown, and the unknowable. PLoS One. 2010 Aug 2;5(8):e11832.

46. Haro-Moreno JM, Rodriguez-Valera F, Rosselli R, Martinez-Hernandez F, Roda-Garcia JJ, Gomez ML, et al. Ecogenomics of the SAR11 clade. Environ Microbiol. 2020 May;22(5):1748–63.

47. Logares R, Rengefors K, Kremp A, Shalchian-Tabrizi K, Boltovskoy A, Tengs T, et al. Phenotypically different microalgal morphospecies with identical ribosomal DNA: a case of rapid adaptive evolution? Microb Ecol. 2007 May;53(4):549–61.

48. Větrovský T, Baldrian P. The Variability of the 16S rRNA Gene in Bacterial Genomes and Its Consequences for Bacterial Community Analyses. PLOS ONE. 2013 Feb 27;8(2):e57923.

49. Duarte CM. Seafaring in the 21St Century: The Malaspina 2010 Circumnavigation Expedition. Limnology and Oceanography Bulletin. 2015 Feb 1;24(1):11–4.

50. Martínez-Pérez AM, Osterholz H, Nieto-Cid M, Álvarez M, Dittmar T, Álvarez-Salgado XA. Molecular composition of dissolved organic matter in the Mediterranean Sea. Limnology and Oceanography. 2017 Nov 1;62(6):2699–712.

51. Ruiz-González C, Logares R, Sebastián M, Mestre M, Rodríguez-Martínez R, Galí M, et al. Higher contribution of globally rare bacterial taxa reflects environmental transitions across the surface ocean. Molecular Ecology. 2019 Apr 1;28(8):1930–45.

52. Pernice MC, Giner CR, Logares R, Perera-Bel J, Acinas SG, Duarte CM, et al. Large variability of bathypelagic microbial eukaryotic communities across the world’s oceans. The ISME Journal. 2016 Apr 1;10(4):945–58.

53. Salazar G, Cornejo-Castillo FM, Benítez-Barrios V, Fraile-Nuez E, Álvarez-Salgado XA, Duarte CM, et al. Global diversity and biogeography of deep-sea pelagic prokaryotes. The ISME Journal. 2016 Mar 1;10(3):596– 608.

54. Sanz-Sáez I. Contribution of marine heterotrophic cultured bacteria to microbial diversity and mercury detoxification. 2021; Available from: http://hdl.handle.net/10261/233620

55. Parada AE, Needham DM, Fuhrman JA. Every base matters: assessing small subunit rRNA primers for marine microbiomes with mock communities, time series and global field samples. Environmental Microbiology. 2016 May 1;18(5):1403–14.

56. Stoeck T, Bass D, Nebel M, Christen R, Jones MDM, Breiner HW, et al. Multiple marker parallel tag environmental DNA sequencing reveals a highly complex eukaryotic community in marine anoxic water. Molecular Ecology. 2010 Mar 1;19(1):21–31.

57. Callahan BJ, McMurdie PJ, Rosen MJ, Han AW, Johnson AJA, Holmes SP. DADA2: High-resolution sample inference from Illumina amplicon data. Nature Methods. 2016;13(7):581–3.

58. Quast C, Pruesse E, Yilmaz P, Gerken J, Schweer T, Yarza P, et al. The SILVA ribosomal RNA gene database project: improved data processing and web-based tools. Nucleic Acids Research. 2012;41(D1):D590–6.

59. Guillou L, Bachar D, Audic S, Bass D, Berney C, Bittner L, et al. The Protist Ribosomal Reference database (PR$^2$): a catalog of unicellular eukaryote Small Sub-Unit rRNA sequences with curated taxonomy. Nucleic Acids Research. 2012;41(D1):D597–604.

60. Sebastián M, Ortega-Retuerta E, Gómez-Consarnau L, Zamanillo M, Álvarez M, Arístegui J, et al. Environmental and physical barriers drive the basin-wide spatial structuring of Mediterranean Sea and adjacent Eastern Atlantic Ocean prokaryotic communities. Submitted. 2021;

61. Boyer TP, Antonov JI, Baranova OK, Garcia HE, Johnson DR, Mishonov AV, et al. World ocean database 2013. National Oceanographic Data Center (U.S.) OCL, editor. 2013; Available from: https://repository.library.noaa.gov/view/noaa/1291

62. Gloor GB, Macklaim JM, Pawlowsky-Glahn V, Egozcue JJ. Microbiome Datasets Are Compositional: And This Is Not Optional. Frontiers in Microbiology. 2017;8:2224.

63. Tackmann J, Rodrigues JFM, von Mering C. Rapid Inference of Direct Interactions in Large-Scale Ecological Networks from Heterogeneous Microbial Sequencing Data. Cell Systems. 2019;9(3):286-296.e8.

64. Deutschmann IM, Lima-Mendez G, Krabberød AK, Raes J, Vallina SM, Faust K, et al. Disentangling environmental effects in microbial association networks. Microbiome. 2021 Nov 26;9(1):232.

65. Bastian M, Heymann S, Jacomy M. Gephi: An Open Source Software for Exploring and Manipulating Networks. ICWSM [Internet]. 2009 Mar 19 [cited 2021 Mar 30];3(1). Available from: https://ojs.aaai.org/index.php/ICWSM/article/view/13937

66. Fruchterman TMJ, Reingold EM. Graph drawing by force-directed placement. Software: Practice and Experience. 1991 Nov 1;21(11):1129–64.

67. Csardi G, Nepusz T. The igraph software package for complex network research. InterJournal. 2006;Complex Systems:1695.

68. Espejo R, Mestre G, Postigo F, Lumbreras S, Ramos A, Huang T, et al. Exploiting graphlet decomposition to explain the structure of complex networks: the GHuST framework. Scientific Reports. 2020;10(1):12884.

69. Pržulj N, Corneil DG, Jurisica I. Modeling interactome: scale-free or geometric? Bioinformatics. 2004;20(18):3508–15.

70. Yaveroğlu ÖN, Malod-Dognin N, Davis D, Levnajic Z, Janjic V, Karapandza R, et al. Revealing the Hidden Language of Complex Networks. Scientific Reports. 2014;4(1):4547.

71. Prim RC. Shortest connection networks and some generalizations. The Bell System Technical Journal. 1957 Nov;36(6):1389–401.

72. McInnes L, Healy J, Saul N, Grossberger L. UMAP: Uniform Manifold Approximation and Projection. The Journal of Open Source Software. 2018;3(29):861.

73. McInnes L, Healy J, Astels S. hdbscan: Hierarchical density based clustering. The Journal of Open Source Software. 2017;2(11):205.

