## Supplementary Material for "Disentangling microbial networks across pelagic zones in the global ocean"

Short title: Marine microbial networks across space

Manuscript for: Science Advances

#### SUPPLEMENTARY MATERIAL

#### SUPPLEMENTARY FIGURES

**Supplementary Figure 1:** Network metrics grouped by region and depth layer.

**Supplementary Figure 2:** Associations occurring in each region and depth layer. If an association appears in more than 20% of subnetworks in each region, it is classified as low-frequency, >50% prevalent, and >70% global. The number of samples appears in the upper left corner, the number of edges in the upper right corner, and the depth range in the lower right corner (in m below surface). We classified the associations considering all six regions (A-D) and considering the five ocean basins without the MS (E-H).

**Supplementary Figure 3:** Regional associations occurring in each region and depth layer. Within a particular depth layer, if an association appears in at least one subnetwork (present) in one region and in no subnetwork (absent) in other regions, it is classified as regional. The four ocean layers (rows) are surface (SRF), DCM, mesopelagic (MES), and bathypelagic (BAT). The number of samples appears in the upper left corner, the number of edges in the upper right corner, and the depth range in the lower right corner (in m below surface).

**Supplementary Figure 4:** ASVs across depth layers. For each region, we color ASVs based on the layer they first appeared: surface (S, yellow), DCM (D, orange), mesopelagic (M, red), and bathypelagic (B, black). Absent ASVs are grouped in box “a”. An ASV only appearing in the bathypelagic, is assigned to box “a” in above layers. That is, an ASV detected in the surface and present in the DCM but absent in lower layers, appears in the box (S) in the surface and DCM layer, but in box “a” in the meso- and bathypelagic layer. An ASV cannot be assigned to two layers. Note that most ASVs in the bathypelagic zone have been already detected in upper layers because most ASVs are assigned to the boxes “S”, “D”, and “M” instead of “B”.

**Supplementary Figure 5:** Robustness of the third condition for generating sample-specific subnetworks for each region and depth with sufficient samples (DCM layer from the SPO was removed because it contained only one sample). Within each region and depth, the set of samples was randomly subsampled containing between 10% to 90% of the original set using all samples. The y-axis shows the fraction of edges that were kept in the subsampled set compared to the original set. We considered A) only the number of kept edges and B) which edges were kept.

### SUPPLEMENTARY TABLES

**Supplementary Table 1:** Fraction of microbial associations across depth layers. For each region and layer (rows), we determined the constitution of associations (in percentage %) classifying them based on their first appearance (columns): surface, DCM, mesopelagic, and bathypelagic. We indicated the fractions above 40% in grey.

| Region | Layer | Surface | DCM | Mesopelagic | Bathypelagic |
|---|---|---|---|---|---|
| MS | SRF | 100.00 | | | |
| | DCM | 45.14 | 54.86 | | |
| | Mesopelagic | 10.35 | 18.42 | 71.24 | |
| | Bathypelagic | 2.73 | 5.12 | 69.71 | 22.44 |
| NAO | SRF | 100.00 | | | |
| | DCM | 68.30 | 31.70 | | |
| | Mesopelagic | 11.64 | 6.59 | 81.77 | |
| | Bathypelagic | 11.62 | 1.35 | 43.49 | 43.54 |
| SAO | SRF | 100.00 | | | |
| | DCM | 45.08 | 54.92 | | |
| | Mesopelagic | 6.15 | 8.50 | 85.35 | |
| | Bathypelagic | 12.22 | 6.30 | 26.97 | 54.61 |
| SPO | SRF | 100.00 | | | |
| | DCM | 50.07 | 49.93 | | |
| | Mesopelagic | 6.44 | 2.66 | 90.90 | |
| | Bathypelagic | 9.81 | 3.32 | 14.15 | 72.71 |
| NPO | SRF | 100.00 | | | |
| | DCM | 54.23 | 45.77 | | |
| | Mesopelagic | 8.33 | 6.06 | 85.61 | |
| | Bathypelagic | 17.46 | 5.34 | 19.92 | 57.28 |
| IO | SRF | 100.00 | | | |
| | DCM | 39.23 | 60.77 | | |
| | Mesopelagic | 5.92 | 7.87 | 86.21 | |
| | Bathypelagic | 11.00 | 3.84 | 29.61 | 55.56 |

**Supplementary Table 2** Subnetwork cluster. Clusters dominated, i.e. over 50%, by one layer or one region are indicated in grey. The last row shows unassigned subnetworks.

| cluster ID | Dominated by | Size | Fraction of depth layers | | | | | Number of regions (if no number if indicated, it is 1x) | | | | |
|---|---|---|---|---|---|---|---|---|---|---|---|---|
| | | | Epipelagic | | | Meso- | Bathy- | Epipelagic | | | Meso- | Bathy- |
| | | | SRF | EPI | DCM | pelagic | pelagic | SRF | EPI | DCM | MES | BAT |
| 1 | MS | 5 | 20.00 | 20.00 | 20.00 | 20.00 | 20.00 | SAO | MS | NAO | MS | MS |
| 2 | MS | 10 | 10.00 | - | 20.00 | 20.00 | 50.00 | MS | - | 2xMS | 2xMS | 5xMS |
| 3 | MS | 8 | 12.50 | - | - | 25.00 | 62.50 | SRF | - | - | 2xMS | 5xMS |
| 4 | MS, MES | 8 | - | 12.50 | - | 75.00 | 12 | - | MS | - | 6xMS | MS |
| 5 | MS, MES | 12 | 16.67 | - | - | 66.67 | 16.67 | IO, NAO | - | - | 7xMS, NAO | 2xNAO |
| 6 | | 8 | 12.50 | 25.00 | 12.50 | 25.00 | 25.00 | IO | MS, NAO | NPO | MS, NAO | 2xMS |
| 7 | BAT | 15 | 13.33 | - | - | 26.67 | 60.00 | IO, SPO | - | - | IO, MS, SAO, SPO | IO, MS, NAO, 2xNPO, 2xSAO, 2xSPO |
| 8 | DCM | 10 | 10.00 | - | 90.00 | - | - | NPO | - | 5xMS, NPO, 3xSAO | - | - |
| 9 | DCM | 11 | 36.36 | - | 63.64 | - | - | 2xNAO, NPO, SAO | - | 3xIO, 2xMS, NPO, SAO | - | - |
| 10 | | 12 | - | - | 8.33 | 50.00 | 41.67 | - | - | NAO | IO, MS, NAO, 2xNPO, SAO | IO, 2xNAO, NPO, SAO |
| 11 | MES | 6 | - | - | - | 83.33 | 16.67 | - | - | - | IO, MS, NPO, 2xSAO | IO |
| 12 | NAO, MES | 6 | 16.67 | - | - | 83.33 | - | NAO | - | - | 2xMS, 3xNAO | - |
| 13 | SRF | 11 | 54.55 | 9.09 | - | 27.27 | 9.09 | IO, MS, NPO, 3xSAO | MS | - | 2xMS, NAO | MS |
| 14 | BAT | 16 | 12.50 | 6.25 | 6.25 | 6.25 | 68.75 | MS, NAO | MS | MS | MS | 5xNAO, 3xNPO, 2xSAO, SPO |
| 15 | SRF | 8 | 100.00 | - | - | - | - | 3xIO, 4xNAO, NPO | - | - | - | - |
| 16 | MS, SRF | 7 | 71.43 | 14.29 | - | 14.29 | - | 4xMS, NPO | MS | - | MS | - |
| 17 | MS | 9 | - | 11.11 | 33.33 | 22.22 | 33.33 | - | MS | MS, NAO, SPO | 2xMS | 3xMS |
| 18 | MS, BAT | 8 | 12.50 | 25.00 | - | - | 62.50 | IO | 2xMS | - | - | 3xMS, 2xNAO |
| 19 | SRF | 7 | 85.72 | 14.29 | - | - | - | 2xIO, NAO, NPO, 2xSAO | MS | - | - | - |
| 20 | SRF | 15 | 73.33 | - | 6.67 | 6.67 | 13.33 | 2xIO, 2xNAO, NPO, 5xSAO, SPO | - | MS | IO | IO, NPO |
| 21 | | 8 | 25.00 | - | 12.50 | 25.00 | 37.50 | IO, SPO | - | MS | MS, SAO | IO, 2xNAO |
| 22 | | 17 | 23.53 | - | 5.88 | 35.29 | 35.29 | 3xSAO, SPO | - | MS | NAO, 2xNPO, SAO, 2xSPO | IO, MS, NAO, 3xSAO |
| 23 | SRF | 8 | 75.00 | 12.50 | - | 12.50 | - | IO, 2xMS, NAO, NPO, SPO | MS | - | MS | - |
| 24 | MS, MES | 13 | 15.38 | 7.69 | - | 61.54 | 15.38 | 2xMS | MS | - | IO, 4xMS, 3xNAO | NAO, NPO |
| 25 | | 14 | 28.57 | 7.14 | 14.29 | 7.14 | 42.86 | 2xMS, 2xNAO | MS | 2xMS | NAO | MS, 3xNPO, 2xSAO |
| 26 | SRF | 7 | 85.72 | 14.29 | - | - | - | 2xIO-SRF, MS-EPI, 2xNAO-SRF, 2xNPO-SRF | 2xIO-SRF, MS-EPI, 2xNAO-SRF, 2xNPO-SRF | - | - | - |
| 27 | SRF | 11 | 100.00 | - | - | - | - | 2xIO, NAO, 4xNP, 4xSPO | - | - | - | - |
| 28 | MS | 11 | 9.09 | 27.27 | - | 36.36 | 27.27 | MS | 3xNAO | - | 4xMS | 3xMS |
| 29 | | 12 | 50.00 | - | 16.67 | 16.67 | 16.67 | IO, MS, 3xNAO, SAO | - | MS, NAO | 2xMS | 2xMS |
| 30 | | 6 | 50.00 | - | 16.67 | 16.67 | 16.67 | IO, NAO, SPO | - | MS | NPO | IO-BAT |
| 31 | MS | 28 | 25.00 | 10.71 | 7.14 | 35.71 | 21.43 | 4xIO, 2xMS, SAO | 3xMS | 2xMS | 6xMS, 2xNAO, 2xNPO | IO, 2xMS, 3xNAO |
| 32 | SRF | 6 | 100.00 | - | - | - | - | IO, 2xNA, NPO, 2xSAO | - | - | - | - |
| 33 | SRF | 6 | 100.00 | - | - | - | - | NAO, 3xNPO, SAO, SPO | - | - | - | - |
| 34 | SRF | 14 | 100.00 | - | - | - | - | IO, 4xNAO, 5xNPO, 2xSAO, 2xSPO | - | - | - | - |
| 35 | SRF | 13 | 69.23 | 7.69 | - | - | 23.08 | 4xIO, 3xNAO, SAO, SPO | MS | - | - | 3xMS |
| 36 | SRF | 7 | 100.00 | - | - | - | - | 3xIO, 3xNPO, SAO | - | - | - | - |
| - | | 24 | 41.67 | - | 12.50 | 29.17 | 16.67 | 2xIO, MS, 2xNAO, 3xNPO, 2xSAO | - | MS, 2xNAO | 2xIO, 4xMS, NPO | MS, NAO, NPO, SAO |

MS – Mediterranean Sea, NAO – North Atlantic Ocean, SAO – South Atlantic Ocean, SPO – South Pacific Ocean, NPO – North Pacific Ocean, IO – Indian Ocean, EPI – epipelagic layer, SRF – surface, DCM – Deep Chlorophyll Maximum, MES – mesopelagic layer, BAT – bathypelagic layer

**Supplementary Table 3:** Number of environmentally-driven edges detected by EnDED. We removed environmentally-driven edges (indirect) from the preliminary network, which contained 31966 edges. Only edges that were not environmentally-driven by any environmental factor (not indirect) remained in the network.

| Environmental factor | Number of samples | indirect | Not indirect |
|---|---|---|---|
| Fluorescence | 394 | 4 (0.01%) | 31962 |
| $NO_3^-$ | 361 | 1563 (4.9%) | 30403 |
| $PO_4^{3-}$ | 359 | 1357 (4.2%) | 30609 |
| Salinity | 395 | 67 (0.2%) | 31899 |
| $SiO_2$ | 360 | 632 (2.0%) | 31334 |
| Temperature | 395 | 622 (1.9%) | 31344 |
| All | | 2848 (8.9%)<br>= 1779 removed by 1<br>+ 751 removed by 2<br>+ 308 removed by 3<br>+ 10 removed by 4 | 29118 (91.1%) |

**Supplementary Table 4** Number of edges within each region and depth layer before (J>0%) and after filtering edges with low Jaccard index measuring how often the association partners appeared together in the region and depth layer. The DCM layer in the South Pacific Ocean (SPO) contained only one subnetwork, which resulted in the edge prevalence being 100% for all edges. To generate the sample-specific subnetworks, we selected the Jaccard index of J>20%.

| Region | Layer | Samples | Depth (m) | J>0% | J>10% | J>20% | J>30% | J>40% | J>50% |
|---|---|---|---|---|---|---|---|---|---|
| MS | EPI - SRF | 19 | 3 | 3710 | 3631 | 3263 | 2881 | 2375 | 1797 |
| | EPI | 18 | 12-50 | 4763 | 4682 | 4196 | 3731 | 3064 | 2189 |
| | EPI - DCM | 21 | 40-130 | 5545 | 5417 | 4736 | 4030 | 3062 | 2027 |
| | MES | 52 | 200-1000 | 8756 | 8403 | 7336 | 6179 | 4629 | 3088 |
| | BAT | 35 | 1100-3300 | 4497 | 4263 | 3694 | 3171 | 2506 | 1830 |
| NAO | EPI - SRF | 34 | 3 | 15862 | 15255 | 13478 | 11449 | 8487 | 5331 |
| | EPI | 4 | 50 | 3027 | 3027 | 3027 | 2778 | 2529 | 2091 |
| | EPI - DCM | 6 | 70-106 | 3865 | 3865 | 3738 | 3480 | 2973 | 2212 |
| | MES | 14 | 200-800 | 6325 | 6289 | 5689 | 5109 | 4169 | 2978 |
| | BAT | 20 | 1200-4539 | 7490 | 7419 | 6831 | 6206 | 5211 | 3857 |
| SAO | EPI - SRF | 26 | 3 | 13118 | 12768 | 11026 | 9269 | 6842 | 4353 |
| | EPI - DCM | 4 | 80-130 | 4199 | 4199 | 4199 | 3941 | 3443 | 2468 |
| | MES | 6 | 450-850 | 3937 | 3937 | 3740 | 3440 | 2687 | 1614 |
| | BAT | 11 | 1290-4000 | 4143 | 4130 | 3886 | 3605 | 3049 | 2254 |
| NPO | EPI - SRF | 29 | 3 | 14376 | 13778 | 11919 | 9907 | 7323 | 4736 |
| | EPI - DCM | 3 | 37-110 | 3100 | 3100 | 3100 | 3100 | 2568 | 1968 |
| | MES | 9 | 200-780 | 4197 | 4197 | 3781 | 3343 | 2583 | 1625 |
| | BAT | 12 | 2000-4000 | 5198 | 5185 | 4834 | 4510 | 4009 | 3372 |
| SPO | EPI - SRF | 14 | 3-5 | 12007 | 11927 | 10420 | 8990 | 6728 | 4480 |
| | EPI - DCM | 1 | 65 | 1530 | 1530 | 1530 | 1530 | 1530 | 1530 |
| | MES | 3 | 450-650 | 2066 | 2066 | 2066 | 2066 | 1756 | 1318 |
| | BAT | 3 | 1500-4000 | 3159 | 3159 | 3159 | 3159 | 2906 | 2128 |
| IO | EPI - SRF | 35 | 3 | 14307 | 13646 | 11736 | 9602 | 6912 | 4396 |
| | EPI - DCM | 3 | 86-130 | 3411 | 3411 | 3411 | 3411 | 2855 | 2310 |
| | MES | 7 | 400-950 | 4654 | 4654 | 4344 | 3961 | 3083 | 2082 |
| | BAT | 8 | 1065-4000 | 2928 | 2928 | 2790 | 2563 | 2101 | 1290 |

MS – Mediterranean Sea, NAO – North Atlantic Ocean, SAO – South Atlantic Ocean, SPO – South Pacific Ocean, NPO – North Pacific Ocean, IO – Indian Ocean, EPI – epipelagic layer, SRF – surface, DCM – Deep Chlorophyll Maximum, MES – mesopelagic layer, BAT – bathypelagic layer

#### SUPPLEMENTARY MATERIAL

**Supplementary Material 1:** Highly prevalent (>70%) associations per region and depth layer. For each association between two taxa (first and second column) we list the frequency (third column) and percentage (forth column) with respect to region (fifth column) and depth layer (sixth column).

**Supplementary Material 2:** Highly prevalent (>70%) regional associations. For each association between two ASVs (first and second column) we list: region (third column), depth layer (fourth column), prevalence in that region and depth layer (fifth column), type: eukaryotic (Euk_Euk), prokaryotic (Prok_Prok), and association between domains (Euk_Prok) (sixth column), and the phyla (seventh and eight column).

**Supplementary Material 3:** Associations appearing in all layers in at least one region. For each association between two ASVs (first and second column) we list: the classification in each layer (3-6 column), overall prevalence (8 column), prevalence in each region and depth layer (9-34 column), the number of regions in which the association appeared in all layers (AllLayers, 35 column), the number of layers an association appears in a region (36-41 column), type: eukaryotic (Euk_Euk), prokaryotic (Prok_Prok), and association between domains (Euk_Prok) (42 column), and the phyla (43-44 column).
