## Supplementary figures and images for "Disentangling microbial networks across pelagic zones in the global ocean"

### Supplementary Figure 1

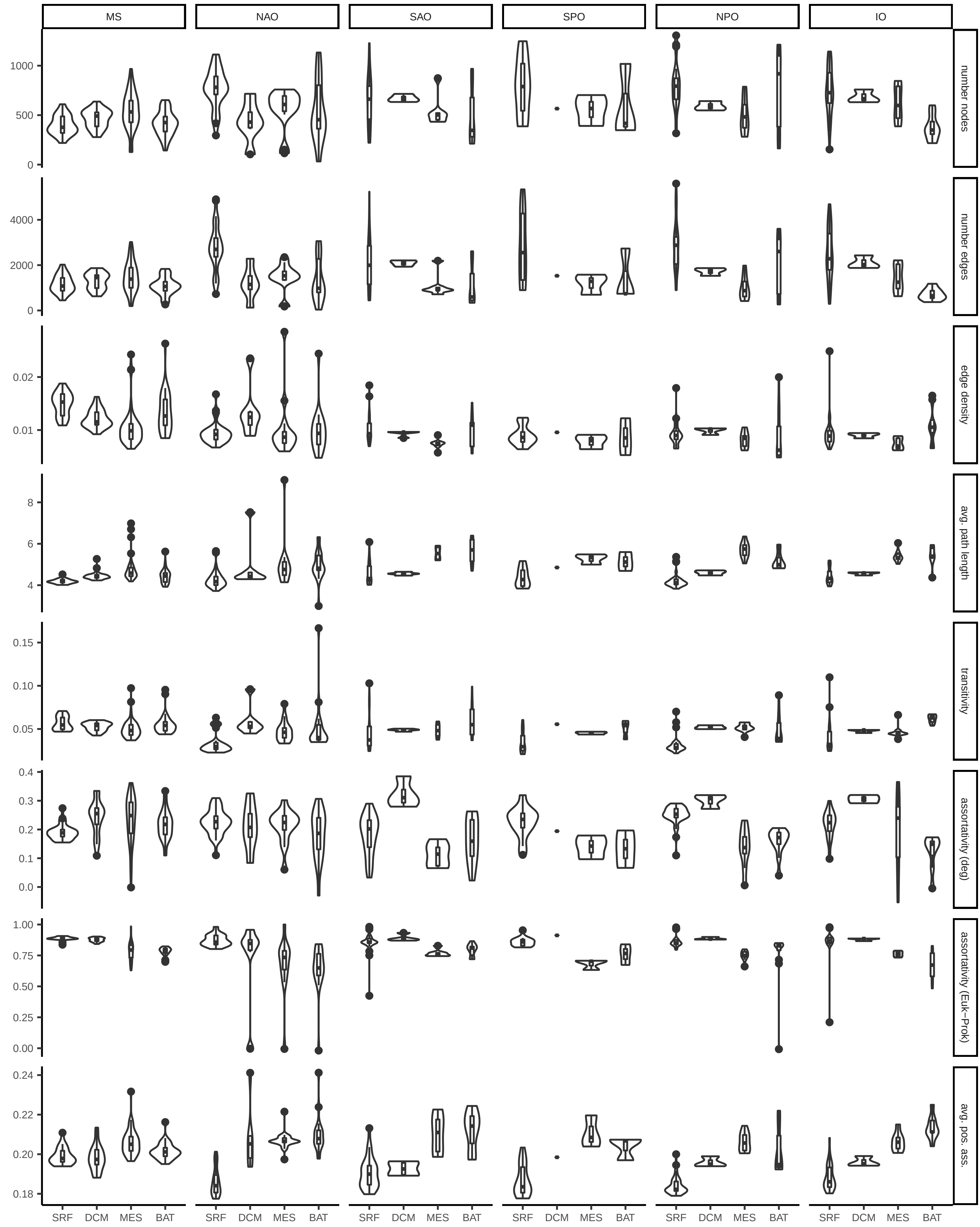

### Supplementary Figure 2

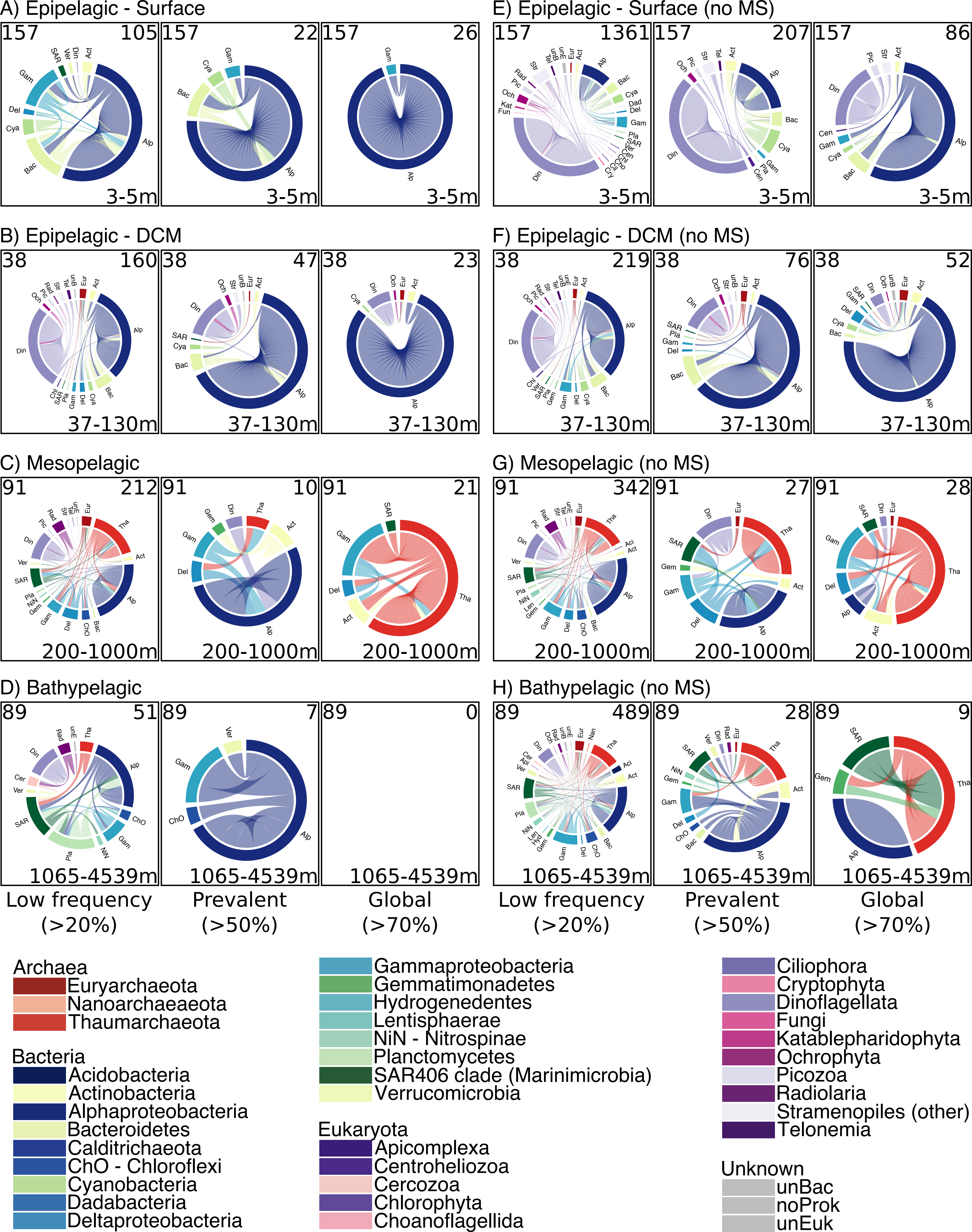

### Supplementary Figure 3

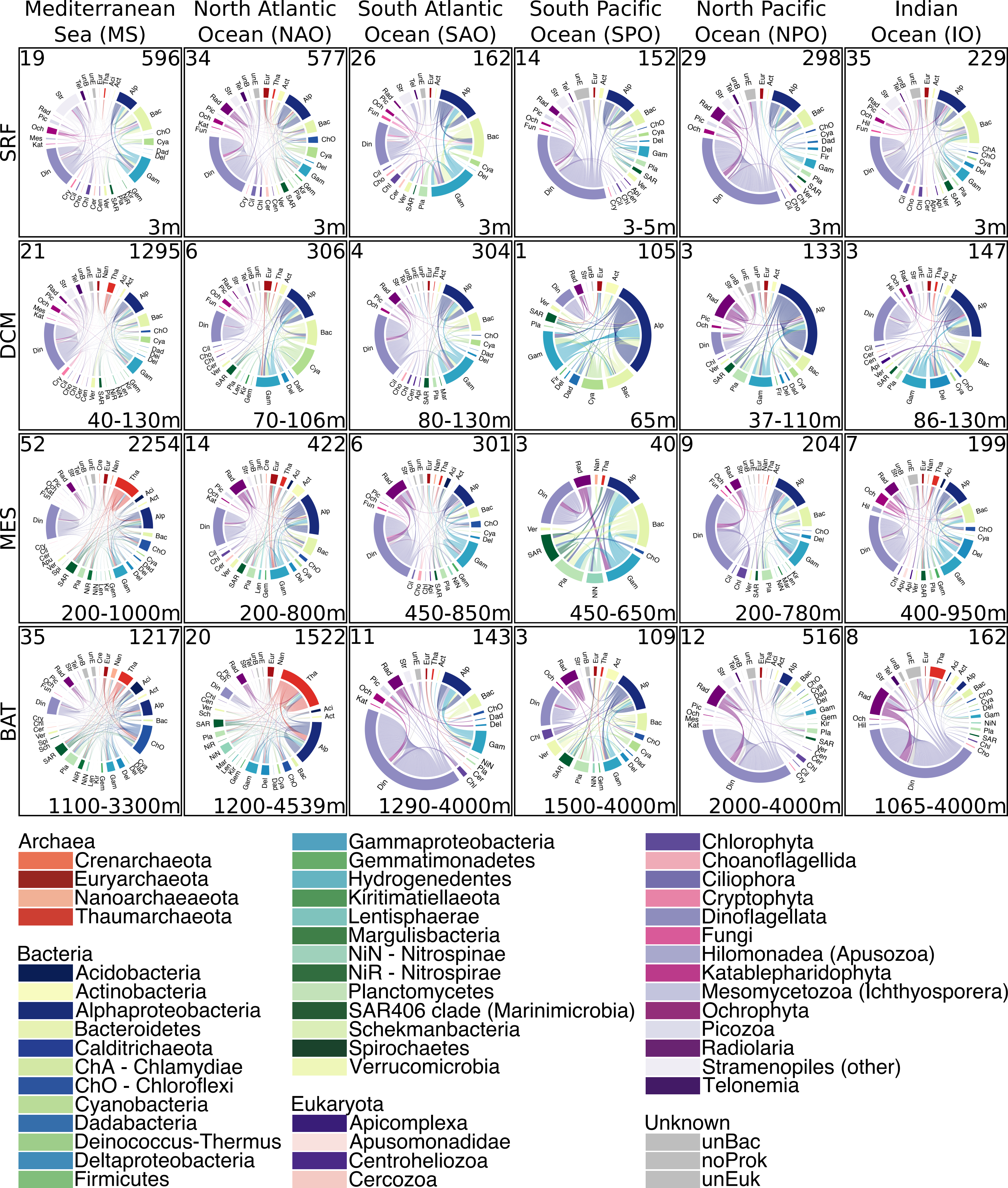

### Supplementary Figure 4

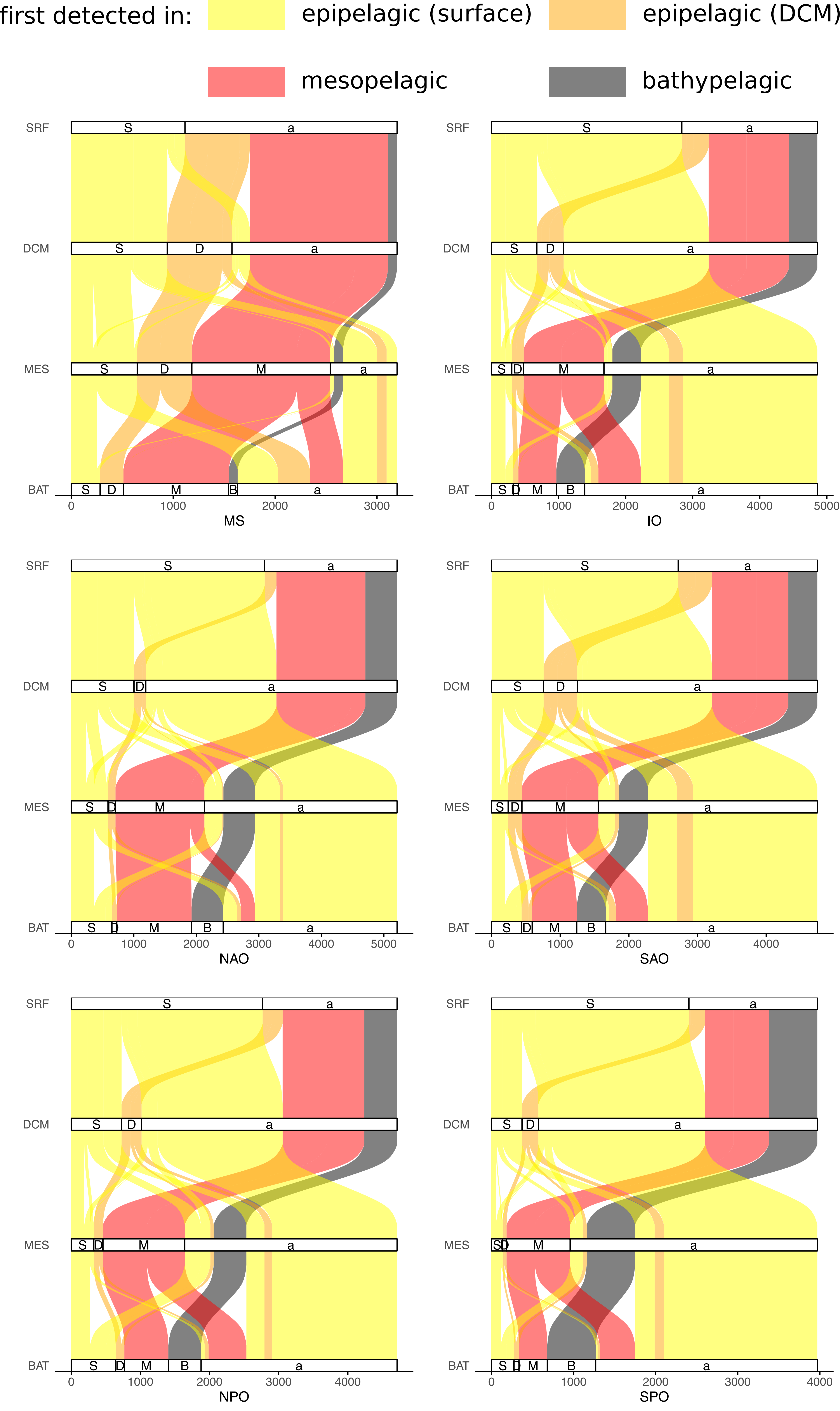

### Supplementary Figure 5

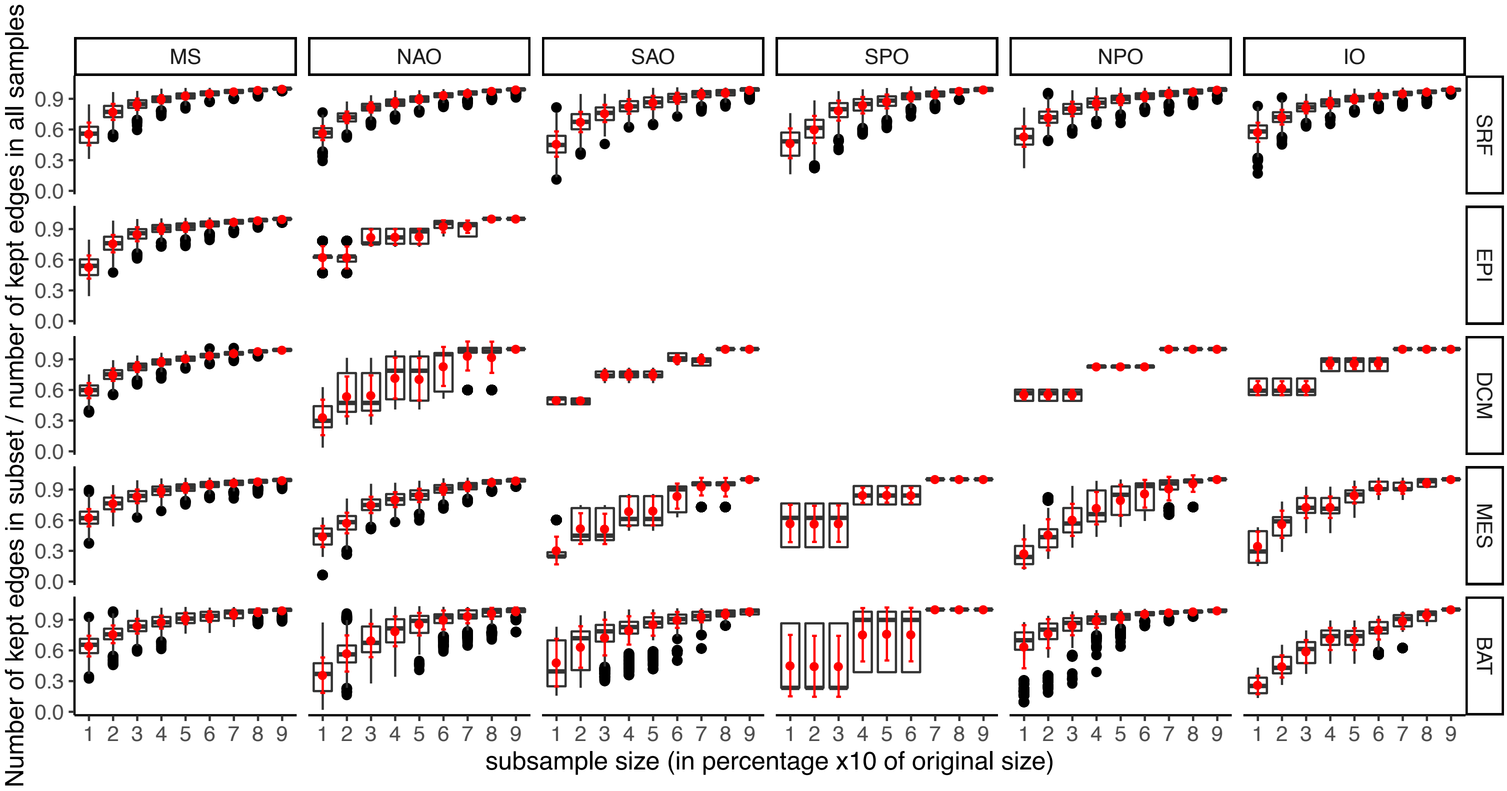

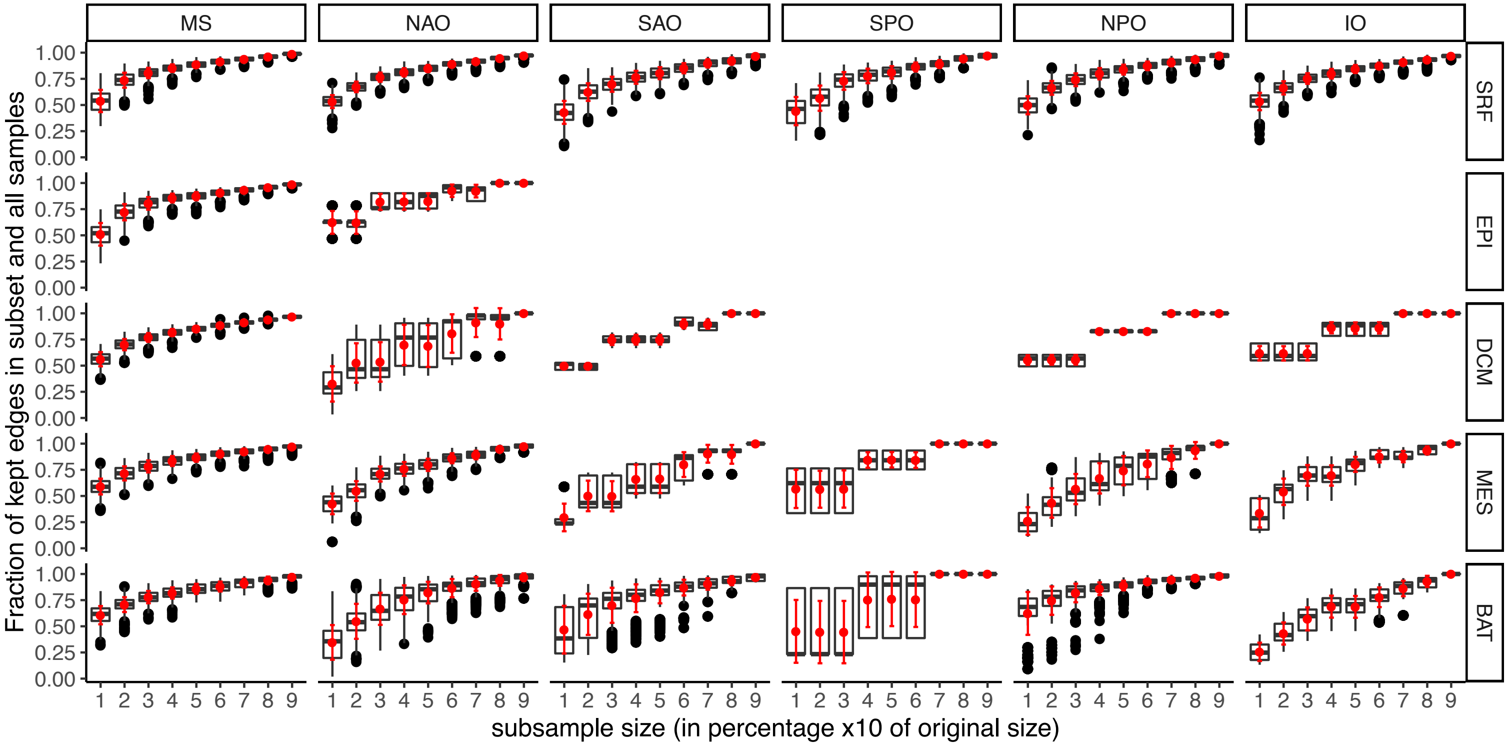
